# Meta-analysis of the genetic loci of pigment pattern evolution in vertebrates

**DOI:** 10.1101/2022.01.01.474697

**Authors:** Joel Elkin, Arnaud Martin, Virginie Courtier-Orgogozo, M. Emília Santos

## Abstract

Vertebrate pigmentation patterns are amongst the best characterised model systems for studying the genetic basis of adaptive evolution. The wealth of available data on the genetic basis for pigmentation evolution allows for meta-analysis of trends and quantitative testing of evolutionary hypotheses. We employed Gephebase, a database of genetic variants associated with natural and domesticated trait variation, to examine trends in how *cis*-regulatory and coding mutations contribute to vertebrate pigmentation phenotypes, as well as factors that favour one mutation type over the other. We found that studies with lower ascertainment bias identified higher proportions of *cis*-regulatory mutations, and that *cis*-regulatory mutations were more common amongst animals harboring a higher number of pigment cell classes. We classified pigmentation traits firstly according to their physiological basis and secondly according to whether they affect colour or pattern, and identified that carotenoid-based pigmentation and variation in pattern boundaries are preferentially associated with *cis*-regulatory change. We also classified genes according to their developmental, cellular, and molecular functions. We found that genes implicated in upstream developmental processes had greater *cis*-regulatory proportions than downstream cellular function genes, and that ligands were associated with higher *cis*-regulatory proportions than their respective receptors. Based on these trends, we discuss future directions for research in vertebrate pigmentation evolution.

## Introduction

One of the central goals of evolutionary biology is to understand the genetic basis of organismal diversity. Determining which genes and mutations underlie adaptive traits is essential for understanding how evolutionary forces shape organismal variation (Barrett and Hoekstra, 2011). In this regard, vertebrate pigmentation is a powerful system for integrating historically disparate fields of evolutionary research, and ultimately identifying the genetic mechanisms underlying evolutionary change (Cuthill et al., 2017; Hubbard et al., 2010; Kronforst et al., 2012; Orteu and Jiggins, 2020).

Vertebrate pigmentation offers a diverse array of phenotypes, with intra- and interspecific variation ranging from whole-body colour changes, to highly localised pattern alterations (Figure 1). Similarly, the adaptive significance of pigment patterns can be attributed to multiple distinct selection pressures, including thermoregulation, camouflage, aposematism, sexual display, and ultraviolet protection (Protas and Patel, 2008). The rapid evolution of vertebrate pigmentation patterns, and their evolutionary significance, allows for both mapping of individual mutations to their resultant phenotypes, and inference of the evolutionary pressures driving the selection of said phenotypes. As such, there have been many studies identifying loci associated with a wide range of pigmentation phenotypes in different vertebrate taxa. A meta-analysis of these individual case studies may reveal broad trends underlying pigment pattern evolution and help to inform the future direction of vertebrate pigmentation research.

**Figure 1.**
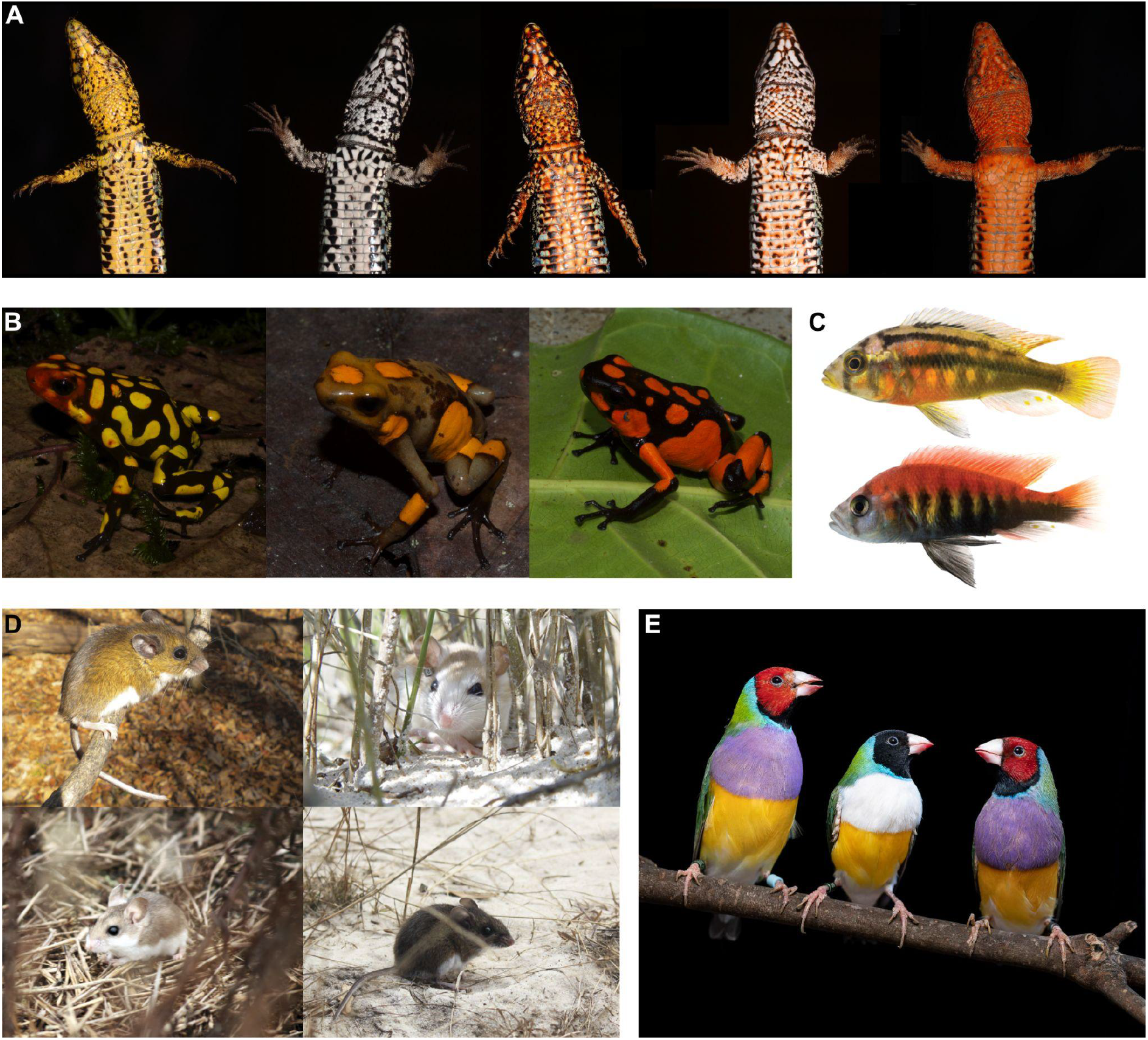
Examples of vertebrate colour pattern diversity. **A:** Sympatric colour morphs of the European common wall lizard (*Podarcis muralis*) (Andrade et al., 2019). The colour morphs differ in a range of key morphological, physiological, or behavioural traits. Photographs courtesy of Pedro Andrade and Miguel Carneiro. **B:** Allopatric morphs of harlequin poison frogs (*Oophaga histrionica* complex). Amphibian colour patterns are extremely under-represented in pigmentation evolutionary genetic studies. Photographs courtesy of Roberto Marquez. **C:** Two species of Lake Victoria cichlids. *Pundamilia nyererei* (top) and *Haplochromis sauvagei* (bottom) showing differences in horizontal stripes and vertical bars (Kratochwil et al., 2018). Photographs courtesy of Claudius Kratochwil. **D:** Different species of *Peromyscus* mice showing differences in colour phenotypes – *P. maniculatus nubiterrae* (top left, image credit to Evan P Kingsley)*, P. polionotus phasma* (top right, image credit to JB Miller)*, P. polionotus sumneri* (bottom left, image credit to Nicole Bedford) *and P. gossypinus* (bottom right, image credit to Nicole Bedford). Differences in melanic coat colouration correlate with the colour of the background substrate and evolved as a result of strong predation. Photographs adapted from Bedford and Hoekstra, 2015. **E:** Red and black head colour polymorphism in the Gouldian finch (*Erythrura gouldiae*), morphs display differences in aggressivity and reproductive success (Toomey et al., 2018). Photograph courtesy of Ricardo Jorge Lopes and Miguel Carneiro.

Here, we first outline the basic biology of vertebrate pigmentation, as well as the key differences between different vertebrate clades. We then utilise a dataset of vertebrate pigmentation literature to analyse trends in the underlying genetics. For this purpose, we used Gephebase (https://www.gephebase.org/) - a knowledge base dedicated to the compilation of literature on genes and mutations underlying natural and domesticated organismal variation in Eukaryotes (Courtier-Orgogozo et al., 2020). Gephebase gathers published data on evolutionarily relevant mutations, each entry representing a causal association between an allelic change at a given locus and trait variation between individuals or species. Each entry includes information relating to the species/population, the type of trait, the gene, the nature of the mutation(s), whether they represent null alleles, and the study that identified it. Using the vertebrate pigmentation entries in this database, we focused on the relative abundance of *cis*-regulatory and coding mutations contributing to vertebrate pigmentation evolution. Finally, we discuss the direction of future vertebrate pigmentation research and the role to be played by recent model systems and innovative approaches.

### Vertebrate pigmentation

In vertebrates, most colour patterns derive from specialised pigment cells which produce either pigments or reflective structures. These cells derive from the migratory neural crest cell population, which emerges from the dorsal neural tube during early vertebrate development (Lapedriza et al., 2014). Neural crest cells then delaminate and undergo some of the longest migrations of any embryonic cell type to give rise to multiple derivatives such as neurons and glia of the peripheral nervous system, smooth muscle, craniofacial cartilage and bone, and pigment cells (Simões-Costa and Bronner, 2015).

Across vertebrate clades there is considerable diversity in the cell types producing colours. In fish, amphibians, and non-avian reptiles, there are multiple distinct classes of pigmented and structurally coloured cells, called chromatophores. These classes contain different combinations of pigments and/or reflective structures, and therefore exhibit different ranges of colours. Across these clades up to at least nine chromatophore classes are recognised. The most common chromatophore classes that utilise pigments are melanophores (brown/black melanin pigments) and xanthophores (yellow/orange carotenoid pigments). These cells produce pigment-based colour via the deposition of their respective pigment molecules, which selectively absorb specific wavelengths of light. In contrast, structural colouration results from the presence of reflective structures, such that structural colour is variable depending on the angle from which it is viewed. The most common structural chromatophore is the iridophore, which appears silvery or blue due to the arrangement of layered purine platelets of variable size, shape, and arrangement (Parichy, 2021; Singh and Nüsslein-Volhard, 2015).

Rarer chromatophore classes include red erythrophores and blue cyanophores, as well as two distinct classes of white leucophores with different regulatory profiles, developmental origins, and chemical compositions (Goda and Fujii, 1995; Huang et al., 2021; Lewis et al., 2019). Each chromatophore class present in an organism goes through extensive cell movements and cell-cell interactions to form a final colour pattern. Variation in the abundance, combinations, and arrangements of chromatophores generates the intricate and diverse colour patterns seen in fish, amphibians, and reptiles.

In contrast, mammals and birds have independently lost most pigment cell diversity and mainly retain only one pigmentary cell type, the melanocyte (equivalent to the melanophore) (Kelsh et al., 2009). Melanocyte differentiation and development remains highly conserved between vertebrates, but differences exist. Unlike melanophores, melanocytes produce two melanin pigment types in different shades – brownish-black eumelanin, and reddish pheomelanin. The ability to switch melanogenesis between eumelanin and pheomelanin production is specific to birds and mammals (McNamara et al., 2021). Mammals and birds develop complex pigmentation patterns by temporally and spatially regulating melanogenesis switching, as opposed to using different chromatophore classes for different colours. Furthermore, while other vertebrates retain pigments within the respective chromatophore, in birds and mammals melanin is secreted from melanocytes into the skin or the integumentary appendages, such as feathers and hairs (McNamara et al., 2021). Finally, birds additionally exhibit an array of yellow, orange and red colours due to the processing and accumulation of dietary carotenoid pigments (Toews et al., 2017).

### The role of *cis*-regulatory and coding mutations in evolution

The extent to which evolution is predictable on the level of genetic variation is a long-standing topic in evolutionary biology. One important question concerns the relative prevalence of *cis*-regulatory and coding sequence mutations (Carroll, 2008; Hoekstra and Coyne, 2007; Martin and Courtier-Orgogozo, 2017; Martin and Orgogozo, 2013; Stern and Orgogozo, 2009, 2008). It has been hypothesised that *cis*-regulatory mutations are more likely to contribute to evolutionary changes in morphology (Carroll, 2008). One reason for this is that *cis*-regulatory mutations are expected to have fewer pleiotropic effects when compared to coding mutations, mostly due to their highly modular nature conferring tissue specificity, affecting gene expression patterns without changing protein function. Conversely, protein coding sequences are less modular, and non-synonymous mutations are expected to affect protein function across every cell and tissue in which it is expressed. For the past three decades though, case studies have shown both types of mutations contributing to evolutionary change. Thus, trying to argue for the existence of a dichotomy in the relative frequency of both types of genetic change is too simplistic. Instead, we should strive to understand when one type of mutation is selected over the other (Stern and Orgogozo, 2008). For example, are different mutations associated with particular aspects of trait variation (e.g. colour versus pattern)? Do the cellular and developmental processes underlying a trait influence the nature of the mutations that can affect it? Do we see a shift in the role played by different types of mutation throughout evolutionary time?

Here, we tackled these questions by examining the relative distribution of *cis*-regulatory and coding mutations across vertebrate pigmentation evolution, and how the relative frequencies of these mutation types associate with other factors, such as loci mapping strategy, type of trait variation or types of genes involved. Moreover, by analysing the types of mutation associated with the evolution of vertebrate pigmentation, we identify trends in the field and discuss possible directions for future research.

## Results

### Genetic variants associated with pigmentation are asymmetrically distributed across vertebrate clades

To examine the representation of different pigmentation systems across vertebrate evolutionary studies, we looked at the number of entries in Gephebase across five clades - teleost fishes, amphibians, squamates, birds and mammals. All of the 363 vertebrate pigmentation entries belonged to one of these clades. The significant majority (71.1%, n = 258) of entries were mammals, whilst birds (20.9%, n = 76) were also highly prevalent (Table 1). Teleosts (5.5%, n = 20) and squamates (2.2%, n = 8) were scarcely represented, and amphibians were associated with only one entry (0.3%). As such, clades in which multiple distinct chromatophore classes are found represent less than 10% of our dataset, in comparison to those clades which harbour only melanocytes and no other pigment cell classes. This taxonomic bias is attributable in part to how the field originated and evolved. For instance, the disproportionate representation of mammalian studies is likely a reflection of the extensive characterisation of the mouse coat colour genetics system, which has been studied for more than a century (Hoekstra, 2006). As a result, many of the key components of the melanin biosynthesis pathway were first identified in mice, and these components are often selected for investigation in other mammalian and bird systems.

**Table 1:**
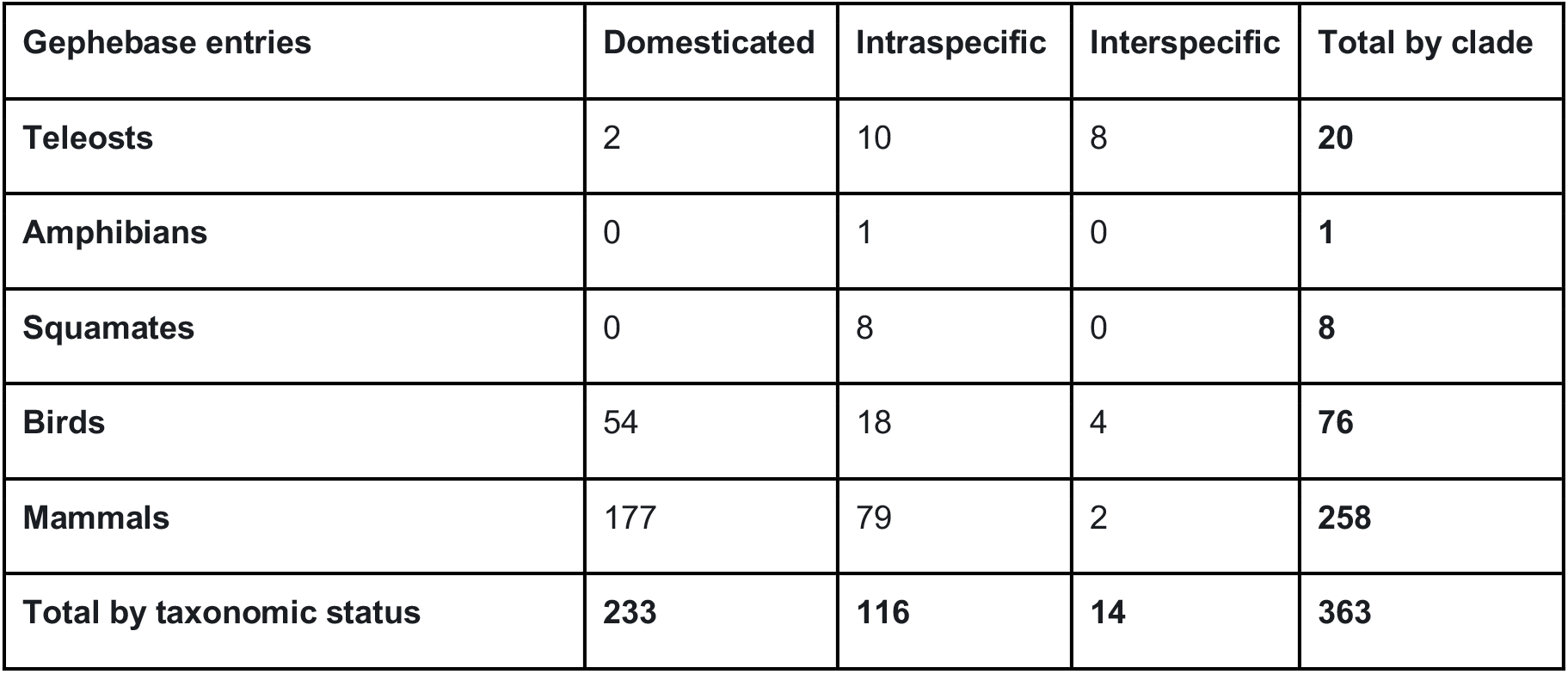
Number of entries in the vertebrate pigmentation Gephebase dataset in 2021 (see methods) according to vertebrate clade and taxonomic status. One entry corresponds to genetic variation at a given gene that has been found to contribute to pigmentation evolution in a given taxon.

| Gephebase entries | Domesticated | Intraspecific | Interspecific | Total by clade |
| --- | --- | --- | --- | --- |
| Teleosts | 2 | 10 | 8 | 20 |
| Amphibians | 0 | 1 | 0 | 1 |
| Squamates | 0 | 8 | 0 | 8 |
| Birds | 54 | 18 | 4 | 76 |
| Mammals | 177 | 79 | 2 | 258 |
| Total by taxonomic status | 233 | 116 | 14 | 363 |

### Variation in vertebrate pigmentation is associated with a majority of coding mutations

We examined the relative prevalence of protein coding and *cis*-regulatory mutations across the dataset. *Cis*-regulatory mutations, and coding sequence mutations together account for 330 entries (90.9%) out of the full dataset, with coding mutations being the majority (262 entries, 72.2%). The remaining classes of genetic variation (9.1%) were gene amplification, gene loss, intronic mutations, and unknown (when the gene has been identified but the exact mutation(s) have not) (Figure 2A). The overall prevalence of coding mutations is in contrast to the hypothesis that *cis*-regulatory mutations have a greater likelihood of generating phenotypic change (Carroll, 2008). This may reflect the relative ease of identifying coding mutations compared with *cis*-regulatory changes (Stern and Orgogozo, 2008). Next we analysed the relationship between the types of mutations and the methodology used to identify them.

**Figure 2.**
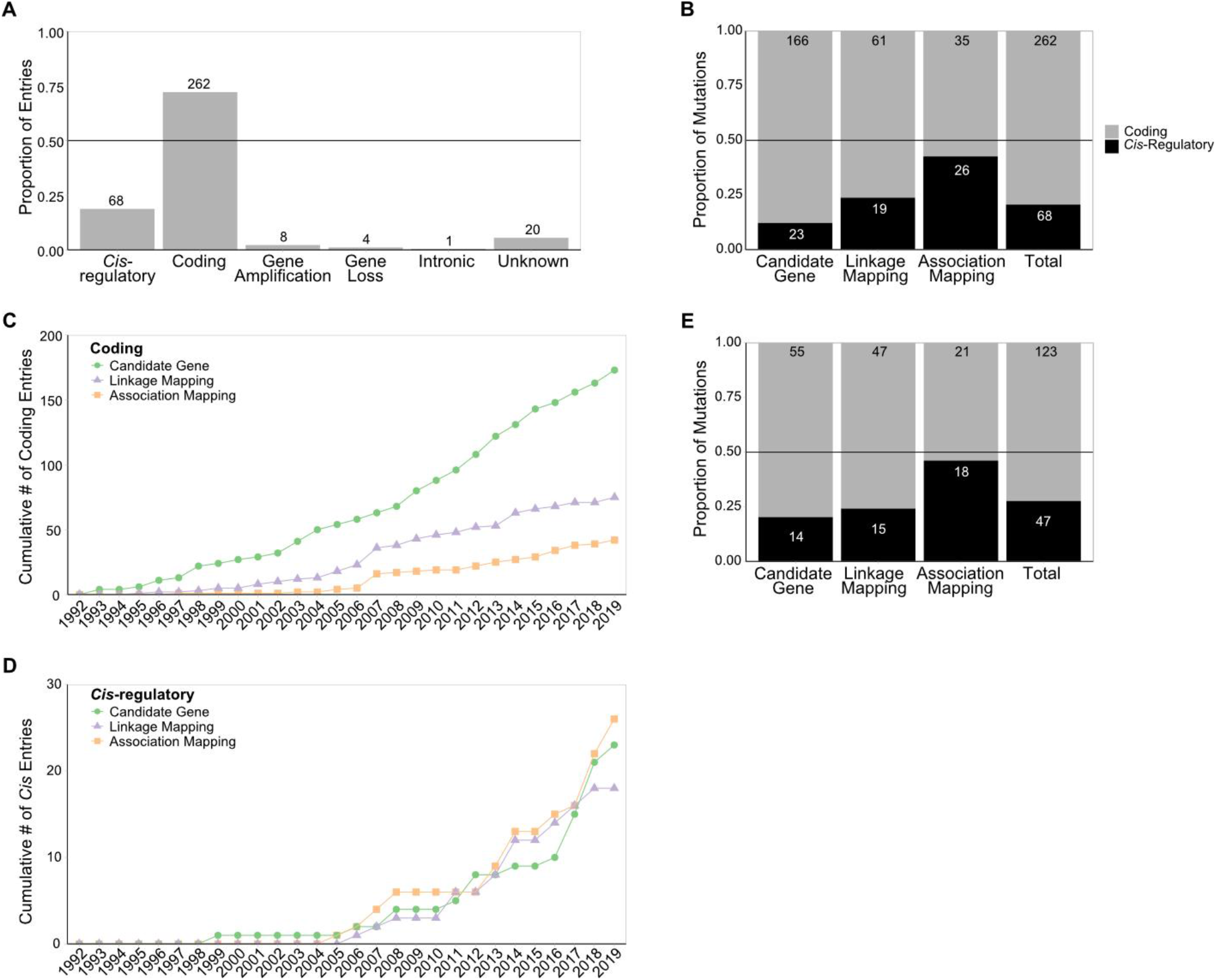
**A:** The relative proportions of all of the mutation types identified. **B:** The *cis*-regulatory and coding mutations associated with each study methodology, as well as the total proportion of the dataset. **C:** The cumulative number of coding mutations of each study methodology over the time period recorded in the Gephebase dataset. **D:** The cumulative number of *cis*-regulatory mutations of each study methodology over the time period recorded in the Gephebase dataset. **E:** The *cis*-regulatory and coding mutations associated with each study methodology, with *mc1r*, *asip* and *kit* entries removed. The numbers above (A) or within (B & E) each bar represent the number of entries in that category. The grey horizontal line is at 0.5.

### Study methodologies with less ascertainment bias exhibit slightly higher proportion of *cis*-regulatory mutations

Three methodology categories are represented in the dataset -– *Candidate Gene*, *Linkage Mapping*, and *Association Mapping* (Courtier-Orgogozo et al., 2020). Candidate gene studies exhibit the highest ascertainment bias and may be expected to identify a higher proportion of coding mutations. Indeed, many candidate gene studies have historically focused on identifying amino-acid changes within coding regions in new species, based on previous findings in other species. However, they also facilitate comparisons across broad taxonomic distances compared with other mapping approaches. In contrast, both linkage and association mapping studies start with a phenotypic difference and attempt to pin it to a sequence change, at least as a locus interval. Linkage mapping involves crossing populations to generate recombinant hybrids, and thus can only map variation between closely related organisms which are interfertile. However, such studies are usually less biased than candidate gene approaches in their detection of causative loci, depending on the resolution and coverage of the mapping. Finally, association mapping approaches involve identification of significant association within large, heavily intermixed populations, and typically have minimal ascertainment bias as a result. To account for potential differences between methodologies we plotted the proportion of *cis*-regulatory mutations for each method (Figure 2B).

As expected, candidate gene approaches identified the lowest proportion of *cis*-regulatory mutations (12.2%) and was the only category below the dataset average. Linkage mapping had a higher proportion (23.8%), and association mapping was higher still (42.6%). All three of these methodology categories reported a higher overall proportion of coding than *cis*-regulatory mutations (Figure 2B). As such, regardless of the method used, protein coding mutations underlie the majority of the investigated vertebrate pigmentation variation. The effect of study methodology on the *cis*-regulatory proportion appears to be that decreased ascertainment bias increases the discovery of causal *cis*-regulatory mutations.

These results have to be taken with caution, since all methodologies will tend to favour the identification of coding mutations. For example, once a candidate region is found by linkage mapping or association mapping, it is still easier to find coding mutations contributing to the trait of interest than *cis*-regulatory mutations: based on the genetic code it is easy to identify mutations disrupting the amino acid sequence of the encoded protein whereas it is difficult to predict *cis*-regulatory effects based on sequence alone. Furthermore, validation of *cis*-regulatory mutations usually requires reporter constructs and transgenic animals, thus being more time-consuming than the validation of coding mutations, which can often involve *in vitro* assays.

### Changes in *cis*-regulatory reporting over time do not fully explain the disparity in mutation types

The Gephebase dataset includes studies published between 1993 and 2020 (Courtier-Orgogozo et al., 2020). Owing to the increasing availability of genomic resources facilitating linkage and association mapping in a broader range of model systems (Kratochwil and Meyer, 2015), we expected the *cis*-regulatory proportion of the dataset to increase over time. We therefore examined the cumulative number of *cis*-regulatory and coding mutations over the time period covered by the dataset, for each study methodology. We used this approach as the low numbers of entries for each individual year made non-cumulative comparison misleading – for instance, there were several years with no entries for a particular study methodology. We found that *cis*-regulatory entries began to appear in the dataset later than coding entries (Figure 2C and 2D). Prior to 2005 only a single *cis*-regulatory mutation was identified, compared with 65 coding. However, past this point *cis*-regulatory entries were added at approximately the same rate as coding, for each study methodology. Thus, although the overall *cis*-regulatory proportion does increase over the entire time period covered, this is largely due to coding mutation discovery beginning earlier.

We also did not observe an appreciable increase in the relative prevalence of linkage mapping or association mapping studies. Taken together, this suggests that the overall trend is a slight increase in *cis*-regulatory mutation reporting, but that differences in the relative prevalence of different study methodologies are not sufficient to explain the disparity between *cis*-regulatory and coding mutations. As such, it is possible that there is a discovery bias towards the identification of coding mutations in all methods, or alternatively, that evolutionary variations in vertebrate pigmentation involve a higher proportion of coding mutations. In the next sections, we further examine this proposition with particular attention to research biases and phenotypic categories.

### The relative frequency of *cis*-regulatory and coding mutations holds in the absence of the most represented genes in the dataset

Historically, most evolutionary studies have centred on the melanic mammalian and bird pigmentation systems owing to the important knowledge in mice coat colour genetics. Therefore, we further explored if the prevalence of protein coding mutations was due to the overrepresentation of melanic pigmentation case studies. Mammal and avian melanic pigment patterns are the result of dynamic switching between the synthesis of two types of melanin. The switch between black eumelanin and reddish-brown pheomelanin is controlled by the melanocortin 1 receptor, MC1R (García-Borrón et al., 2005). *Mc1r* is the most represented gene in our dataset, with 87 entries that account for 26.4% of the dataset. The second-most represented gene in the dataset is *kit*, a cell-surface receptor that is essential for the expression of tyrosinase, a rate-limiting enzyme in melanin biosynthesis (Hou et al., 2000). 40 *kit* entries account for 12.1% of entries. Finally, the third-most represented gene is agouti signalling protein, *asip*. ASIP acts as an antagonist to MC1R - in the absence of ASIP, MC1R initiates a signalling cascade resulting in eumelanin production, while ASIP antagonism reverts the melanocyte to pheomelanin synthesis (García-Borrón et al., 2005). There are 33 *asip* entries, corresponding to 10% of the dataset.

The high representation of these three genes in Gephebase reflects their relevance in mammalian and other vertebrate systems, and particularly their frequent selection as candidate genes for further study. To remove a potential bias of methodology resulting from a focus on melanic genes, we performed the same analysis with these three entries - *mc1r*, *kit,* and *asip* - removed. When removing *mc1r*, *kit*, and *asip* entries from the dataset, the overall proportion of *cis*-regulatory mutations increases from 20.6% to 27.6%, owing to the prevalence of coding mutations amongst *mc1r* entries in particular (Figure 2E). The *cis*-regulatory proportion of all three methdology categories increases, and association mapping remains the category with the highest *cis*-regulatory proportion, followed by linkage mapping (Figure 2E). Still, coding mutations remain the majority of entries for each type of experimental evidence. Taken together, the overrepresentation of these melanic case studies does not explain the higher prevalence of protein coding mutations in the vertebrate pigmentation dataset.

### The proportion of coding mutations is higher for studies of domesticated and intraspecific variation than for interspecific variation

It has been previously hypothesised that different kinds of mutation occur with different frequencies during short-term and long term evolution with coding mutations being more common across short-term evolution, and thus shorter taxonomic distances (Stern and Orgogozo, 2009, 2008). Over shorter periods of time, mutations with deleterious pleiotropic effects might be selected for if alternative, less pleiotropic mutations do not immediately appear. Whereas, over long periods of time, non-optimal mutations will be tested in a variety of environments and there will be more opportunity for adaptive mutations without pleiotropic effects to appear and be selected for. In addition, contexts of artificial selection and favourable breeding conditions by humans can overcome the cost of mutations that would normally be counter-selected in the wild (Cieslak et al., 2011; Courtier-Orgogozo and Martin, 2020; Hanly et al., 2021). To test if the same is true for vertebrate pigmentation systems, we plotted the proportion of *cis*-regulatory mutations by taxonomic status. The dataset includes three categories of taxonomic status - *Domesticated*, *Intraspecific*, and *Interspecific*. Gephebase also includes ‘*Intergeneric or higher*’, but no vertebrate colouration entries were assigned this category. The domesticated category included cases of artificial selection by breeders and fanciers, with pigment trait variations being directly selected in most cases. The intraspecific category contained studies that investigated natural phenotypic differences between morphs of the same species. Finally, interspecific entries were all those that used pairs of taxa above the species level.

Domesticated and intraspecific variation show a similarly low proportion of *cis*-regulatory mutations (16.6% and 23.1%, respectively; Figure 3A, Figure S1). In contrast, interspecific variation showed a notably higher proportion of *cis*-regulatory mutations (72.7%), suggesting that their prevalence may increase with increasing taxonomic distance (Figure 3A). This result has to be taken with caution as the number of studies addressing vertebrate pigmentation variation between species is extremely low – only a total of 11 entries were interspecific. Further, 6 out of these 11 case studies were teleost fish entries, which have multiple pigment cell types which could influence the mutation proportion (Figure S2). The low number of interspecific studies together with low clade diversity, limits the examination of trends across large evolutionary timescales and may reflect an underexplored aspect of vertebrate pigmentation evolution.

**Figure 3.**
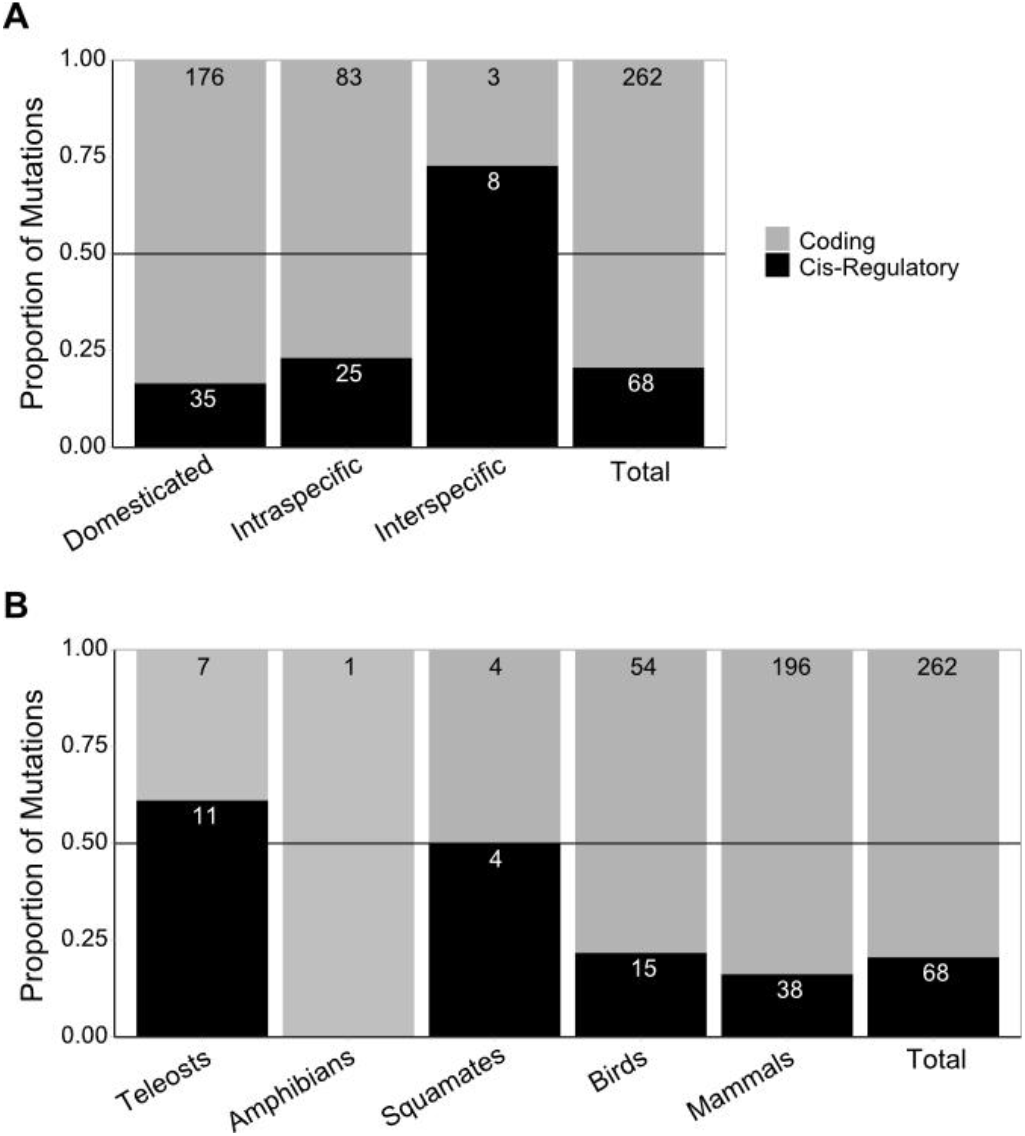
**A:** The proportion of *cis*-regulatory and coding mutations associated with each taxonomic status, as well as the total proportion for the dataset. **B:** The proportion of *cis*-regulatory and coding mutations associated with each clade. The numbers within each bar represent the number of entries in that category. The grey horizontal line is at 0.5.

### The proportion of coding mutations is higher in mammal and bird studies than in teleosts, amphibians, and squamates studies

The prevalence of coding mutations could be associated with the disparity in representation of different vertebrate clades. Birds and in particular mammals constitute the significant majority of entries (91.8%, n = 303 combined), with teleosts, amphibians, and squamates being much rarer (8.2%, n = 27 combined). These clades have divergent pigmentation systems, and notably only teleosts, amphibians, and squamates utilise multiple developmentally distinct classes of pigment cells. This may cause differences in evolutionary trends. For instance, the development of distinct pigment cell classes from a shared neural crest origin increases the likelihood of coding sequence mutations, in genes contributing to pigment cell development, having pleiotropic effects . Similarly, genes implicated in direct cellular interaction might be expected to suffer negative effects from coding sequence mutations, owing to the specificity of recognition that is typical of signalling molecules and their receptors. We therefore examined whether clades with distinct pigmentation systems lead to differences in the mutation proportion. For this purpose, we plotted the proportion of mutations by the five vertebrate clades (Table 1, Figure 3B). The only amphibian entry in the dataset was coding. Excluding amphibians, mammals had the lowest proportion of *cis*-regulatory changes (16.2%). Birds had a similarly low *cis*-regulatory proportion (21.7%). In contrast, squamates and teleosts both showed higher rates of *cis*-regulatory change - 50%, and 55.0%, respectively.

Although the low number of entries associated with multiple chromatophore pigment systems does preclude drawing definitive conclusions, this trend fits with the hypothesis that variation in pigmentation systems utilising multiple pigment cell classes exhibit a higher proportion of *cis*-regulatory mutations. Teleosts, amphibians, and squamates combined had a *cis*-regulatory proportions of 55.5%, compared with 17.5% for birds and mammals combined. Notably, teleosts exhibited a much higher proportion of interspecific studies than the dataset average - 6 of the 18 teleost entries were interspecific (Figure S2). The strong association between teleost entries and interspecific comparison makes it difficult to conclude which (if either) is the more important factor in yielding higher *cis*-regulatory proportions. Although, when taking into account only the intraspecific cases, the proportion of *cis*-regulatory mutations is still higher in teleosts, amphibians, and squamates (42.1%, 8 out of 19) than in birds and mammals (23.6%, 17 out of 72) (Figure S2). A larger sample of studies conducted in model systems harbouring multiple pigment cell classes, together with more interspecific studies in other clades, would provide valuable insight in this regard.

### Types of phenotypic variation associated with coding versus *cis-*regulatory mechanisms

Differences in the nature of phenotypic variations may be associated with distinct types of genotypic change. For instance, loss of pigmentation may be more likely to result from loss-of-function coding sequence changes, whereas patterning changes could be preferentially driven by *cis*-regulatory mutations due to their high spatial modularity reducing pleiotropic effects (Orteu and Jiggins, 2020). To determine whether there are differences between colour loss, colour tuning, and pattern variation, we assigned different phenotype categories to each entry (Table 2).

**Table 2:** Summary of phenotype categories: assignment criteria for each phenotype category, and the total number of entries in the dataset for which that category was assigned.

| Category Name | Criteria for Assignment | Number of Entries | Example |
| --- | --- | --- | --- |
| **Colour loss versus colour tuning** |  |  |  |
| Pheomelanism | A phenotype exhibiting increased abundance of pheomelanin relative to eumelanin, through changes to melanogenic switching or production of either melanin type. This includes phenotypes in which eumelanin synthesis is absent or reduced such that pheomelanin is relatively more prevalent. | 61 | A missense mutation in *MC1R* is associated with the red phenotype in domesticated donkeys (Abitbol et al., 2014). |
| Eumelanism | A phenotype exhibiting increased abundance of eumelanin relative to pheomelanin, through changes to melanogenic switching or production of either melanin type. This includes phenotypes in which pheomelanin synthesis is absent or reduced such that eumelanin is relatively more prevalent. | 71 | A loss-of-function SNP in *ASIP* is associated with 'black panther' phenotype melanistic leopards (Schneider et al., 2012). |
| Dilution | A phenotype in which both eumelanin and pheomelanin synthesis is reduced (but not lost) across one or several tissues. | 20 | A melanophilin splice site SNP leads to exon skipping and coat colour dilution in domesticated rabbits (Lehner et al., 2013). |
| Amelanism | A phenotype characterised by a loss of melanin production across one or several tissues. Includes conditions variably defined as albinism and leucism. | 42 | Three separate *tyrosinase* coding mutations are associated with oculocutaneous albinism in three species of frog (Miura et al., 2017). |
| Loss of carotenoid coloration | A phenotype resulting from the inability to uptake or process dietary carotenoids causing total loss of carotenoid-based pigmentation. | 2 | A splice site mutation in *SCARB1* inhibits cellular carotenoid uptake, leading to a total loss of carotenoid pigmentation in white canaries (Toomey et al., 2017). |
| Change in carotenoid coloration | A phenotype resulting from changes to the relative or overall abundance of carotenoid-based pigments in the body. | 6 | Gain of expression of a carotenoid ketolase, *CYP2J19*, is responsible for processing yellow carotenoids to produce red ketocarotenoids in red canaries (Lopes et al., 2016). |
| White spotting | A phenotype resulting from impairment of migration and/or differentiation of melanocytes or their precursors, causing patchy regions of white tissue. Differentiated from amelanism through molecular mechanism rather than visual appearance. | 52 | White spotting phenotypes in domestic horses are associated with a large number of distinct mutations identified in *KIT* (Haase et al., 2015). |
| **Colour shift versus pattern alteration** |  |  |  |
| Colour shift | Any phenotype in which organismal colour shifts either locally or globally, without impacting the spatial boundaries of its existing patterning. Includes phenotypes in which an organism's existing pattern is overridden, e.g. whole body eumelanism. | 164 | In Japanese quail, a frameshift deletion in *ASIP* is associated with the recessive black phenotype, in which feather colour is darkened across the whole body without altering pattern boundaries (Hiragaki et al., 2008). |
| Pattern alteration | Any phenotype in which the spatial boundaries of an organism's colour pattern change, including the formation of new boundaries e.g. establishment of a pattern in a previously uniform region. Does not include phenotypes in which an existing pattern changes colour without change to the pattern boundary. | 99 | The convergent evolution of horizontal melanic stripes in different cichlid species is associated with regulatory changes in *agouti-related peptide 2*, and expression levels are predictive of stripe presence across multiple cichlid radiations (Kratochwil et al., 2018). |

Firstly, one of seven categories was assigned reflecting the nature of the phenotype in terms of the distribution of colour across the body - pheomelanism, eumelanism, dilution, amelanism, white spotting, carotenoid change, or carotenoid loss (Table 2). All of these categories encompassed melanic traits except for carotenoid change and loss. Each entry was assigned to one of these categories on the basis of both the visible phenotype as well as the molecular mechanism involved. For example, a phenotype involving patchy regions of white pigmentation could be assigned to amelanism or white spotting, depending on whether pigment synthesis or pigment cell migration was impaired.

Secondly, we separately assigned each entry to colour shift or pattern alteration, on the basis of whether that phenotype represented a shift in colour or a change to spatial pattern boundaries (Table 2).

In every category except changes in carotenoid pigmentation, coding sequence changes were the most prevalent (Figure 4A). The categories with the lowest *cis*-regulatory proportion were loss of carotenoid colouration (0%), and amelanism (4.8%). Although loss of carotenoid colouration had too few entries (only 2) to draw definitive conclusions, it is notable that both categories represent a loss of pigmentation. In contrast, the other five categories represent either changes in relative pigment quantity or impairment of cell migration. The prevalence of coding mutations amongst phenotypes involving pigment loss is largely due to loss-of-function mutations affecting genes vital for carotenoid processing or melanin biosynthesis, respectively. Thus, these results support the hypothesis that phenotypes involving changes to pigmentation quantity or distribution are associated with a higher proportion of *cis*-regulatory mutations than loss phenotypes.

**Figure 4.**
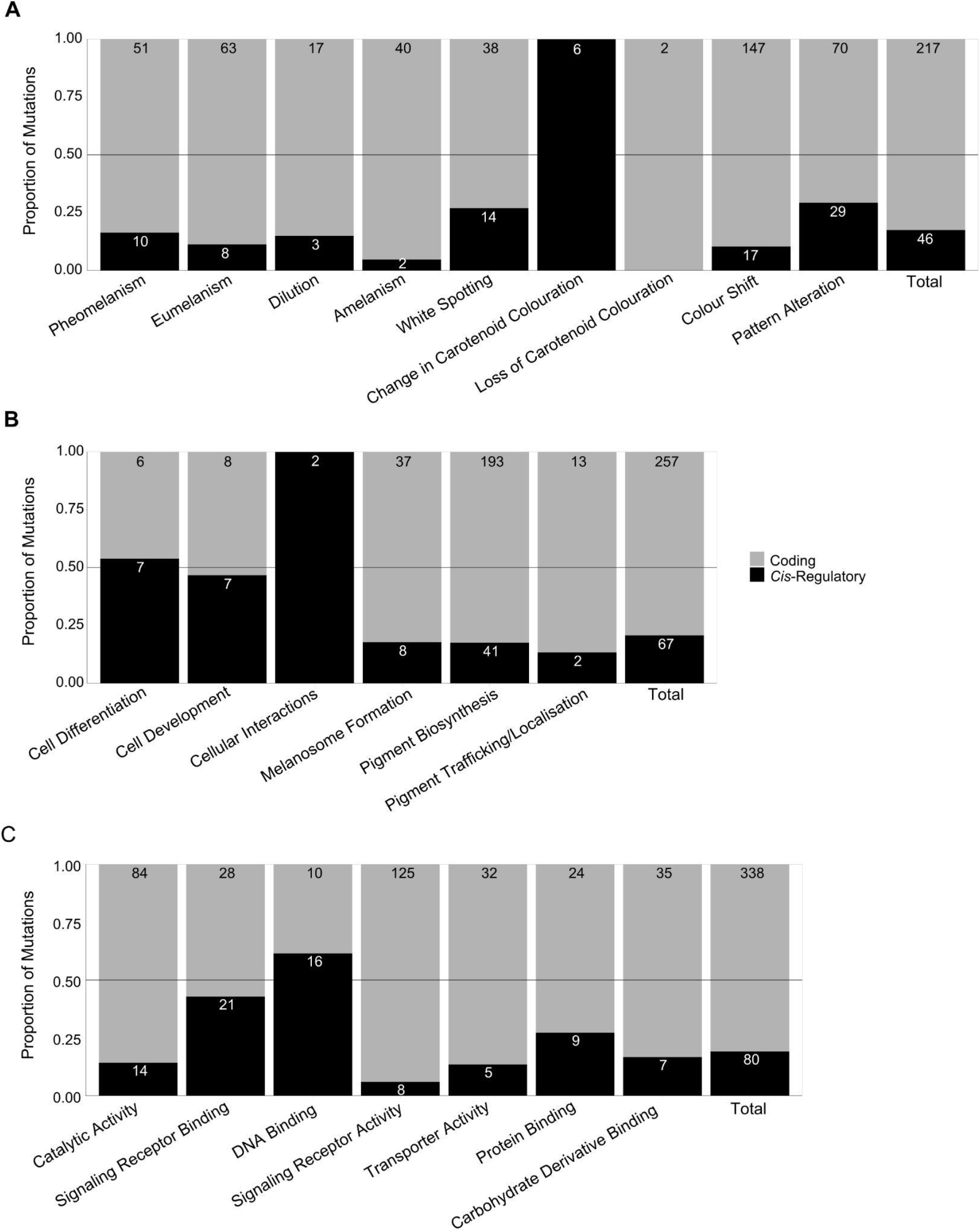
**A:** The proportion of *cis*-regulatory and coding mutations associated with each phenotype category, as well as the total proportions of the dataset. Note that the total proportions were for all entries that were assigned to either colour shift or pattern alteration. **B:** The proportions of *cis*-regulatory and coding mutations associated with each cellular function category. **C:** The proportion of *cis*-regulatory and coding mutations associated with each parent gene ontology assignment. The numbers within each bar represent the number of entries in that category. The grey horizontal line is at 0.5.

Both phenotype categories pertaining to carotenoid colouration had relatively few entries - six for change in carotenoid colouration (all of which were *cis*-regulatory) and two for loss of carotenoid colouration. However, both categories combined did have a high *cis*-regulatory proportion (75.0%) suggesting that carotenoid pigmentation evolution may be more driven by *cis*-regulatory changes compared with melanic pigmentation. The overall *cis*-regulatory proportion for melanic pigmentation was 17.7%. Further studies investigating carotenoid phenotypes would be invaluable in determining whether carotenoid colouration does evolve through different mutation types.

For melanic pigmentation, the category with the highest *cis*-regulatory proportion was white spotting (26.9%), and this was the only melanin-related category with a higher proportion than the average. Amelanism (4.8%) had the lowest proportion, and the remaining categories - pheomelanism (16.4%), dilution (15.0%), and eumelanism (11.3%) were all similar to the average. Notably, two phenotype categories were strongly associated with domesticated entries - 18 out of 20 dilution entries, and all of the 52 white spotting entries, were identified in domesticated taxa (Figure S3). White spotting phenotypes are associated with highly pleiotropic effects, with many white spotting entries in Gephebase being deleterious in the homozygous state. It is therefore not surprising that white spotting mutations were only identified in domesticated taxa, where they can be subject to heterozygous advantage (Hanly et al., 2021; Hedrick, 2015). The slightly higher *cis*-regulatory proportion of this category may reflect the nature of white spotting phenotypes, which result from impairment of pigment cell migration or differentiation. It is therefore possible that coding mutations that would cause white spotting are more likely to be non-viable due to their pleiotropic effects on upstream cellular development.

Finally, phenotypes relating to changes in pigment pattern boundaries had a higher *cis*-regulatory proportion (29.3%) than phenotypes relating to colour shifts (10.4%). This was also true for every study methodology and taxonomic status (Figure S3 and S4). However, notably this difference between colour and pattern phenotypes was more pronounced in non-domesticated studies - for intraspecific and interspecific entries pattern phenotypes had a *cis*-regulatory proportion of 60% compared with 14.3% for colour (Figure S3). This is in spite of white spotting phenotypes being exclusively domesticated as well as exclusively categorised as pattern alterations. Thus, the disparity between colour and pattern phenotypes in terms of mutation type may be larger than this dataset suggests in non-domesticated evolution, particularly given the overall prevalence of domesticated studies in pigmentation research. Overall the Gephebase dataset supports a disproportionate role of *cis*-regulatory changes in the generation of pattern variation (Orteu and Jiggins, 2020).

### Upstream developmental processes are associated with a higher proportion of *cis*-regulatory mutations

Differences in mutation proportions may also be associated with the developmental or cellular function fulfilled by the gene in question. We hypothesise that genes associated with upstream developmental processes would exhibit different proportions to those associated with downstream processes. For example, genes that play a role in cellular differentiation (upstream process) may be less tolerant of coding sequence mutation than genes contributing only to pigment deposition (downstream process). To test this, we assigned each gene to only one of six categories reflecting different cellular and/or developmental processes (Table 3). Three of these categories are cell type-specific functions such as melanosome formation, pigment biosynthesis and pigment trafficking/localisation. The remaining categories included more upstream functions such as pigment cell development, differentiation and cellular interactions (Table 3).

**Table 3:** Summary of functional categories: assignment criteria for each functional category, and the total number of entries in the dataset for which that category was assigned. Each gene was assigned to only one functional category.

| Category Name | Criteria for Assignment | Number of Unique Genes | Number of Entries |
| --- | --- | --- | --- |
| Pigment cell differentiation | Contributes to determining fate specification of pigment cell precursors | 8 | 15 |
| Pigment cell development | Necessary for full development of pigment cells post-specification | 10 | 24 |
| Cellular interactions | Involved in signalling or other cellular interactions, either between or within pigment cell classes | 2 | 2 |
| Melanosome formation | Contributes to the formation or maintenance of melanosomes, specialised melanin vesicles | 8 | 46 |
| Pigment biosynthesis | Role in directly synthesising or processing pigments or their direct precursors | 25 | 253 |
| Pigment trafficking / localisation | Role in inter- or intracellular localisation of pigments, whether as a direct transporter or as a trafficking protein | 4 | 16 |

As predicted, the three categories representing more upstream cellular/developmental processes - pigment cell differentiation, pigment cell development and cellular interactions - were all associated with a higher proportion of *cis*-regulatory mutations than categories representing downstream cellular functions (Figure 4B). Only 2 entries fit the criteria for the cellular interactions category, both of which implicate *cis*-regulatory changes (Figure 4B). This small number of entries makes it difficult to determine whether differences in cellular interactions are associated with different mutation proportions - more data entries would be needed for conclusions to be drawn. Alternatively, the categories representing downstream processes - melanosome formation, pigment biosynthesis and pigment trafficking/localization - all exhibited a higher proportion of protein coding mutations. The overall trend appears to fit the hypothesis that for upstream processes such as pigment cell lineage specification, mutations are more likely to be *cis*-regulatory when compared with downstream cellular processes which do not affect cell viability, such as variable rates of pigment deposition.

### DNA binding proteins are associated with a higher proportion of *cis*-regulatory mutations than other molecular functions

We were also interested in whether the specific molecular activity of a gene would influence the distribution of mutation types. Coding mutations are constrained by the degree of functional conservation required by the protein’s molecular function. Transcription factors implicated in multiple regulatory functions may be more constrained than enzymes that catalyse pigment-specific pathways, as even loss-of-function mutations may be tolerated in enzymes that are functionally specialised and non-essential (Pál et al., 2006). For example, differential expression of the transcription factor *sox10* resulting from *cis*-regulatory mutations underlies changes to melanogenesis in both pigeons and chickens (Domyan et al., 2014; Gunnarsson et al., 2011). *Sox10*, and other *sox* family transcription factors, are highly pleiotropic and essential for neural crest specification, migration and differentiation (Sarkar and Hochedlinger, 2013). No *sox10* coding mutations were identified in this dataset. Conversely, multiple presumptive null coding mutations have been identified in *mc1r*, including one example of parallel evolution in cavefish. Thus, in contrast to *sox10* and other transcription factors, highly specialised proteins such as *mc1r* may be less susceptible to pleiotropy arising from null mutations. As such, we expected that molecular functions that are less pleiotropic would exhibit higher *cis*-regulatory proportions, and vice versa.

Using EBI’s QuickGO mouse slimmer and manual combination of highly related molecular GO terms, we assigned each entry to one of 7 categories of GO molecular function (see Methods, Table S1, Figure S4A and S4B). We then examined the mutation proportions for each GO category (Figure 4C). DNA binding had the highest *cis*-regulatory proportion of any category (61.5%) and was the only category with a majority of *cis*-regulatory mutations. Signaling receptor binding had a higher than average *cis*-regulatory proportion of 42.9%, but conversely signaling receptor activity had the lowest with 6.0% . Protein binding also exhibited a higher than average *cis*-regulatory proportion (27.3%). Taken together, these results indicate that the ability of Transcription factors to evolve phenotypically-causal coding variants is limited by their pleiotropic roles in development, while their non-coding regions offer a more viable path for phenotypic changes.

An interesting result is that ligands (“signaling receptor binding”) shows a higher *cis*-regulatory proportion than receptors (“signaling receptor activity”). For most signaling receptor genes, the corresponding ligand was also present in the dataset in the signaling receptor binding GO category. This includes *kit* and its ligand, as well as multiple endothelin signaling receptors and their respective ligands. Our results suggest that the coding sequence of genes encoding signaling ligands are more constrained than that of their signaling receptors. This supports previous findings that *cis*-regulatory mutations in ligand genes drive morphological evolution (Martin and Courtier-Orgogozo, 2017). The receptor/ligand pairing with the most entries in the dataset, *mc1r/asip*, supports this - *mc1r* has a *cis*-regulatory proportion of 1.1%, in contrast with its ligand *asip* with 39.4%. This trend is also observed when candidate gene studies are removed, with 5.5% of *cis*-regulatory mutations for *mc1r* and 50% of *cis*-regulatory changes for *asip*. A similar example, albeit with fewer entries, is *kit/kitlg*. *Kit* has a cis-regulatory proportion of 17.5% (7 out of 40, 23% if candidate gene studies are excluded), and multiple putative null mutations. Although *kitlg* has only four entries, all uncovered by linkage mapping or association mapping, three are *cis*-regulatory. Both the low number of *kitlg* entries and its relatively higher *cis*-regulatory proportion may indicate a lower evolutionary tolerance of coding mutations when compared with its receptor. Overall, our results indicate that ligands may be more vulnerable to pleiotropy than their corresponding receptors, or that ligand mutations preferentially drive localised pigmentation evolution compared with receptor mutations. It has been suggested that ligands play a specific role in altering spatially localised phenotypes due to the specificity of their expression patterns - in contrast to their corresponding receptors, which are more likely to be ubiquitously expressed (Martin and Courtier-Orgogozo, 2017). Additionally, it may be expected that only the ligand-binding domains of receptor proteins are highly constrained by specificity, where other functional domains are more able to tolerate mutations (Worth et al., 2009). Taken together, both the modularity associated with ligand expression patterns as well as the greater number of mutation-tolerant domains in receptors may explain the higher *cis*-regulatory proportion in ligand encoding genes compared to receptor genes.

### Divergent pigmentation systems have fewer shared evolutionary hotspots

One of the major outcomes of the study of the genetic basis of evolutionary change is the discovery that the repeated co-option of the same genes underlies the evolution of convergent phenotypes. Genes that are repeatedly found as causal drivers of similar phenotypic changes in diverse taxa or populations have been referred to as genetic hotspots (Martin and Orgogozo, 2013). Here, we hypothesised that similar pigmentation systems would be more likely to evolve through re-use of the same set of genes, whereas more divergent pigmentation systems would have fewer shared evolutionary hotspots. We therefore divided the dataset by clade and compared the genes shared between each pairing of clades. The overlap between clades was calculated by combining the entries corresponding to each shared gene, and normalising by the total number of entries in the respective clade (see Methods).

When comparing pigmentation systems with multiple pigment cell classes (teleosts and squamates) to those with only one (birds and mammals), each clade exhibited the greatest hotspot overlap with the clade with which it shares a pigmentation system (Figure 5). The two largest normalised values were in mammalian entries for genes shared with birds, and in bird entries for genes shared with mammals. Likewise both teleosts and squamates had the greatest overlap with one another. Additionally, there was considerable overlap between squamates and birds which may reflect their phylogenetic relationship. As the number of shared hotspot genes would be expected to reduce over evolutionary time, phylogenetic distance is likely to contribute to the observed overlap between clades (Conte et al., 2012).

**Figure 5.**
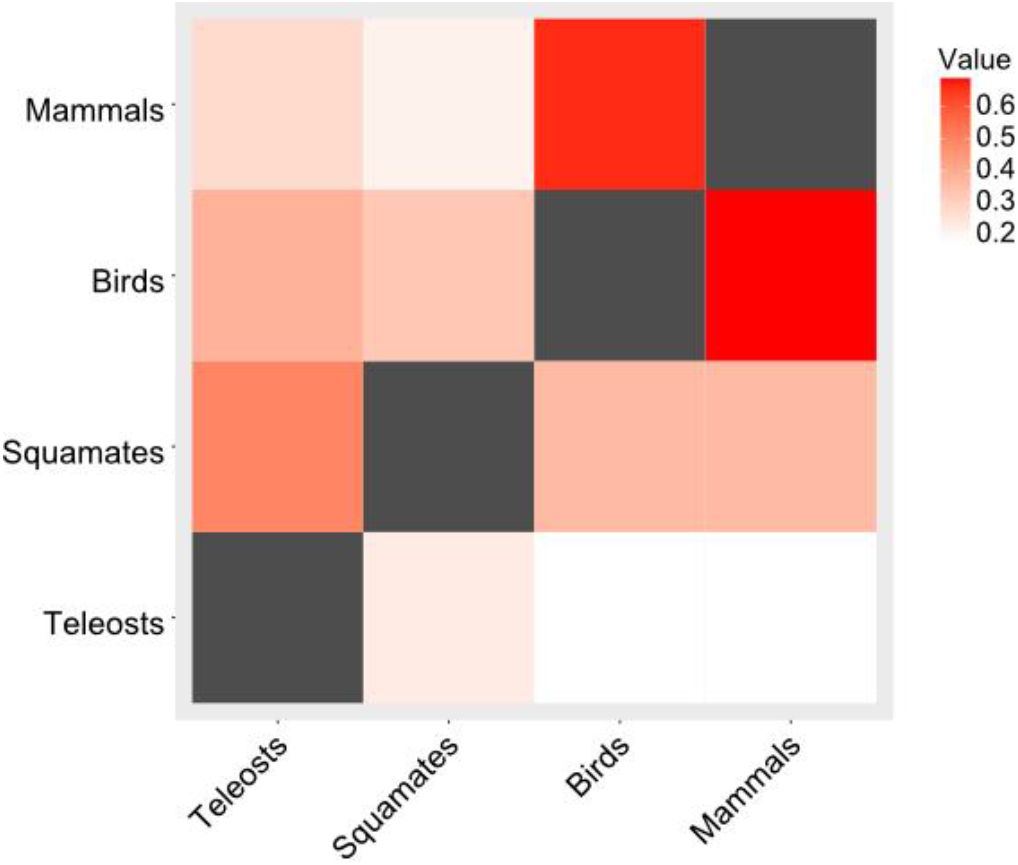
Heatmap of the normalised entries associated with genes shared between each clade pairing. For instance, the top left square has a value equal to the number of mammalian entries for genes that were also identified in teleost studies, divided by the total number of mammalian entries.

Although *mc1r* was highly represented in all clades, and thus contributed considerable hotspot overlap to all clade pairings, many other genes that were highly represented in the dataset were confined to mammals and birds only. The second, third, and fourth most represented genes in the overall dataset - *kit*, *asip*, and *tyrp1* - all had no teleost or squamate entries, and are all implicated in melanogenesis or melanogenic switching. However, certain melanogenic genes also exhibited overlap between teleost and squamate entries, such as *oca2*, a melanosomal transport protein (Table S2). *Oca2* comprised 15% (n = 3) of all teleost entries, and 12.5% (n = 1) of squamate entries - in contrast being rare in mammals (0.82%, n = 2) and absent entirely in birds. Only one non-melanic pigmentation gene was shared between birds and squamates. Aside from *mc1r,* no single gene was identified in all four clades.

Overall, the observed trend of higher hotspot overlap between the most similar pigmentation systems was as expected, and may indicate shared genetic mechanisms underlying pigment evolution in systems with or without multiple pigment cell classes. The low number of non-melanic pigmentation entries (only 22 in total) makes it difficult to draw conclusions about the relevance of hotspot genes outside of melanogenic pathways for clades with additional pigment types.

## Discussion

Gephebase is a powerful tool that can be used to compile data on phenotype-genotype associations to infer general patterns regarding the mechanisms underlying evolutionary change. Importantly, this dataset can be used to find knowledge gaps in the current literature, and identify novel directions and study systems for future research. Here, we analysed a large dataset of 363 Gephebase entries, each a pair of alleles linked to phenotypic variation in vertebrate pigmentation. Of the three forms of empirical evidence accepted by Gephebase, it was clear that the study methodology with higher ascertainment bias - candidate gene approach - identified more coding sequence mutations than the other methods. Irrespective of this bias, protein coding changes were still the majority of mutations identified with linkage and association mapping. In addition, we did identify several factors that may affect the relative distribution of coding and *cis*-regulatory mutations. Below we discuss these factors and highlight new avenues of research in the vertebrate pigmentation field.

### Strong selection pressure might explain the higher proportion of coding mutations associated with vertebrate pigmentation diversity

We found that vertebrate pigmentation overall, and in domesticated species in particular, is associated with a majority of coding mutations. In the case of domesticated vertebrates, this may be explained by the strong selective pressure of the domestication process. During any form of artificial selection, desirable traits will be under strong directional selection, which can overcome the negative pleiotropic effects of coding mutations. Such an example is that of frame overo horses, where changes in patterning are a result of heterozygosity for a coding mutation in endothelin B receptor (*ednrb*), which leads to a white spotted pattern (Metallinos et al., 1998). Homozygosity for this mutation leads to lethal white foal syndrome - affected foals are completely white, and die shortly after birth. Selection for white spotted patterns in domestic horses has thus led to a lethal deleterious coding mutation becoming prevalent in the population, whereas in wild populations such a mutation would likely not have conferred sufficient advantage to overcome its negative pleiotropy.

A sufficiently strong selection pressure may likewise allow the emergence of pleiotropic coding mutations in wild populations. Gephebase contains examples that demonstrate how rapidly changes in pigmentation patterns may evolve when under such pressures, particularly in the context of retaining cryptic camouflage against a new or rapidly changing environment. For instance, populations of North American deer mice exhibit high gene flow, and thus low population structure, with the exception of the agouti locus. Strong selection for crypsis against variable soil colouration instead leads to high variance in agouti coding sequence, and rapid divergence driven by colonisation over the last ∼4000 years (Pfeifer et al., 2018). The prevalence of intraspecific coding mutations in our dataset may therefore be related to the strength of selection typical for vertebrate pigmentation evolution. In the future it would be interesting to investigate the effect of relative selection strength on the role of each mutation type, and to perform the same analyses in multiple morphological traits to determine if these trends are pigmentation specific.

### Evolutionary timescales and taxonomic distance

We found that shorter taxonomic distances (domesticated and intraspecific entries) are associated with a higher proportion of protein coding changes than interspecific changes. These results should be taken with caution as the sparsity of interspecific studies indicates comparatively little investigation of pigmentation traits across larger evolutionary timescales. Interestingly, the majority (8 out of 14) of interspecific entries were identified in teleost species, despite their overall underrepresentation in the dataset. Further, 6 of these interspecific entries were found in East African cichlids. The current overrepresentation of shorter evolutionary timescales may introduce bias in the types of mutation that can be identified. Shorter evolutionary timescales may allow for pleiotropic mutations, as well as mutations that confer an advantage only in a specific environmental context. As such, the higher number of intraspecific and domesticated case studies is likely to result in overrepresentation of coding mutations. Expanding the timescales investigated in pigmentation evolution will facilitate testing of this hypothesis, and a greater understanding of how pigmentation changes over evolutionary time.

Identifying bird and mammal model systems for interspecific studies is challenging owing to the often-limited potential for hybridisation, restricting the types of mapping techniques available. For instance, the only two interspecific mammalian entries examined differences within the black rat species complex, and between the house mouse and tobacco mouse, respectively (Kambe et al., 2011; Robbins et al., 1993). Both of these studies used a candidate gene approach. In birds, Toews et al. used an association mapping approach to examine pigmentation evolution in two naturally hybridising species of warbler with highly divergent pigment patterns (Toews et al., 2016). Diversification of model systems may facilitate a greater understanding of long-term pigment evolution in birds and mammals especially.

### Potential biases introduced by study systems

Mammalian and avian systems dominate the majority of the entries. These only possess melanocytes as pigment cells, and pigmentation differences primarily result from spatial and temporal differences in pigment production and deposition. In other vertebrates, pigmentation patterns result from the presence and differential arrangement of different classes of pigment cells that share a developmental origin. Those few studies in the Gephebase dataset that focus on clades with distinct chromatophore classes often found novel pigmentation phenotypes associated with a wider range of genes. This is also corroborated by the greater numbers of shared hotspot genes between clades with similar pigmentation systems - the genes identified in teleost and squamate studies showed little overlap with those found in mammals and birds. Studies in clades with distinct chromatophore classes also implicated more regulatory mutations than average. As current research continues to contribute new case studies, it will be interesting to look into whether this trend is borne out, or whether the higher prevalence of *cis-*regulatory mutations is an artefact of a low number of studies focusing on organisms with multiple pigment cell classes. In either case, greater emphasis on a broader set of model systems will benefit our broad understanding of vertebrate pigmentation evolution.

In particular it would be of interest to expand the number of studies in teleosts and reptiles to validate the hypothesis that pigmentation systems with multiple chromatophore classes evolve preferentially through *cis*-regulatory modifications. Similarly, amphibian pigmentation evolution was only represented by one study. Amphibians’ highly permeable skin and lack of epidermal protection, as well as their commonly biphasic aquatic and terrestrial life cycles, may necessitate distinct adaptive functions for colouration compared with other vertebrate clades (Rudh and Qvarnström, 2013). These underrepresented clades will be highly informative, particularly with respect to interactions between chromatophore classes and pattern formation.

### The importance of understanding the cellular and developmental basis of variation

We found that changes in patterning had a higher *cis*-regulatory proportion than changes in colour. This is consistent with previous hypotheses that *cis*-regulatory mutations largely govern spatial changes due to altering expression patterns (Carroll, 2008). For example, in sticklebacks a *cis*-regulatory allele of the kit ligand gene reduces its specific expression in gill tissue, leading to divergent gill pigmentation phenotypes (Miller et al., 2007). Gephebase gathers several such examples, where mutations in widely expressed genes are *cis*-regulatory, so that only the expression in specific tissues is affected.

Nonetheless, the majority of pattern alteration case studies involve coding mutations. When looking more closely at some of these, we realised that many are in fact loss-of-function mutations in genes that are expressed only in specific regions of the body. For instance, the urucum breed of canary exhibits a red bill and legs, which are yellow in wild type canaries (Gazda et al., 2020). This is the result of a mutation that compromises the enzymatic activity of BCO2, an enzyme responsible for breakdown of full-length red carotenoids into shorter apocarotenoids. The specificity of this allele in affecting the bill and the legs is the result of loss-of-function mutations in a gene whose expression was already restricted to specific tissues.

Without knowing the expression patterns and the level of pleiotropy of a given gene, we may be wrongly categorising phenotypes as pattern alterations when in fact they are colour shifts. Unfortunately, we have no way of testing this, since most case studies focus solely on the genotype-phenotype association without conducting gene expression or developmental studies. Knowledge of a gene’s specific function, together with its placement in a cellular and developmental context, is essential for an understanding of when and how certain genes and mutations are favoured over others.

Characterising trait development at the gene expression and cellular level is ever more important since we found different mutation proportions associated with differences in the upstream or downstream position of cellular and developmental processes. The genes contributing to upstream processes, i.e. those related to cellular differentiation and development, were more likely to be *cis*-regulatory compared with downstream processes. A previous study (Stern and Orgogozo 2008 Evolution, Table 3) distinguished two categories of genes, the ones that act at or near the terminal points of regulatory networks, named differentiation gene batteries, and the ones that are upstream in the gene network, and found that the proportion of *cis*-regulatory mutations was not significantly different for the two categories. This analysis was based on a compilation of 331 mutations for all types of traits across animals and plants. It is possible that this previous categorization was erroneous as it encompassed very diverse gene networks. Here, by focusing on vertebrate pigmentation, we performed a more robust assignment of the upstream versus downstream gene network position and uncovered an effect on the proportion of *cis*-regulatory mutations.

One of these studies identified a *cis*-regulatory mutation that altered the expression of colony stimulating factor 1 (*csf1*), a signalling factor expressed in xanthophores that is required for their differentiation and survival (Patterson et al., 2014). In *Danio albolineatus,* a close relative of the zebrafish *D. rerio*, increased expression via a *cis*-regulatory mutation of *csf1a* was associated with early xanthophore recruitment (Patterson et al., 2014). The precocious xanthophore appearance led to changes in abundance and distribution of all three principal chromatophore classes (melanophores, xanthophores, and iridophores) which ultimately inhibited formation of the stripe pattern found in other *Danio* species. This study illustrates the potential complexity of phenotypes arising from mutations that affect cellular interactions. The relative lack of studies that have identified such mutations represents an exciting avenue for future pigmentation evolution research.

In contrast, coding mutations in more downstream categories would typically be expected to have lower impact on non-pigmentation phenotype. One study found that a loss-of-function mutation in scavenger receptor B1 (*SCARB1)* in canaries produces a completely white phenotype due to *SCARB1* being necessary for carotenoid uptake. As a result of *SCARB1’*s specialised role, this null coding mutation produces a spectacular pigmentation phenotype (Toomey et al., 2017). These results support the hypothesis that developmentally upstream genes may be constrained by pleiotropy. Given the overrepresentation of pigment synthesis case studies in the vertebrate pigmentation dataset, the collection of more data regarding other types of trait variation is imperative.

### What are the main knowledge gaps?

Most of the vertebrate pigmentation entries (341 out of 363) result from the study of variation in melanic traits. In contrast, variation in other types of colour traits, such as carotenoids and pteridines, structural colouration, and colour plasticity remain largely unexplored. Further, most case studies address naturally selected traits with a relatively simple genetic basis – and only 20 out of 363 entries assessed variable sexually dimorphic traits. It would therefore be interesting to test if the same findings hold true for sexually selected traits or highly polygenic traits where each mutation has a small effect size.

The benefits of broadening the nature of the focal trait are manifold, but an obvious one is that by focusing on unexplored traits we may find previously uncharacterised genes underlying these traits. For example, the study of variation in yellow pigmentation in budgerigar parrots led to the identification of a coding mutation in a previously uncharacterised polyketide synthase that is involved in the synthesis of yellow polyene pigments called psittacofulvins (Cooke et al., 2017).

We found only one entry in the dataset relating to cellular structural colouration, an interspecific study investigating pigment pattern differences between two *Danio* fish species. *D. nigrofasciatus* has a disrupted stripe pattern in comparison to *D. rerio*, which is associated with a *cis*-regulatory mutation in the *Endothelin-3* (*Edn3*) signaling peptide (Spiewak et al., 2018). *Edn3* promotes iridophore proliferation, this *cis*-mutation leads to decreased *Edn3* expression and correspondingly lower iridophore complement in *D. nigrofasciatus*. A lower number of iridophores secondarily affects melanophore proliferation, leading to a reduction of both cell types and disruption of the defined striped pattern in *D. nigrofasciatus.* This study being the only one in Gephebase relating to structural variation highlights that structural colouration is relatively unexplored, and represents an emerging field of research where new genes and developmental processes leading to variation in cells and physical structures will be identified.

The current entries in Gephebase are heavily focused on colouration traits that are genetically determined independently of environmental variables. Colour variation can also arise as a result of phenotypic plasticity, where the same genotype will generate different colour states under different conditions. Six entries pertained to plastic or seasonal colour changes – for instance, *cis*-regulatory alleles of *Asip* determine whether the winter coat of snowshoe hares is brown or white (Jones et al., 2018). Determining how plasticity emerges and evolves remains a challenge. Genes involved in plasticity evolution can only be identified when mapping differences between closely related species or populations which show variation in the presence and absence of plastic colouration (Gibert, 2017). Likewise variation in plastic colour responses can only be mapped by measuring variation in populational reaction norms, which are costly and time consuming experiments. Nonetheless, such studies will be key to discern between the two main hypotheses regarding the evolution of plasticity – whether evolutionary change occurs through changes in genes that sense and regulate a downstream response to external factors, or instead through changes in the colouration genes themselves that become responsive to environmental cues.

Sexually selected traits are also underrepresented in the database, making up only 5.5 % of the case studies. Understanding the interplay between natural and sexual selection on colour traits is important, since they may often act in opposite directions. For example, in several species of Lake Malawi cichlids a well camouflaged colour morph is associated with a *cis*-regulatory mutation in *pax7*. However, this mutation also has a deleterious effect in that it disrupts male nuptial colouration. This conflict has been resolved by the invasion of a novel sex determination locus in tight linkage with the *pax7* allele (Roberts et al., 2009). The importance of pigmentation patterns in mate choice can lead to such conflicts, and the ways in which they are resolved can present fascinating case studies. Sexually selected traits also present a conundrum in that it is unclear why and how trait variation is maintained in natural populations despite apparent directional selection due female choice or male-male competition (Chenoweth and McGuigan, 2010). Identifying genes and genomic regions contributing to trait variation and studying its adaptive significance in wild populations creates the opportunity to understand this paradox (Johnston et al., 2013).

Finally, the Gephebase dataset is currently biased towards large-effect loci, as linkage mapping is limited in its statistical power to detect small effect mutations and polygenic architectures (Rockman, 2012; Slate, 2013). Association studies conducted on thousands of samples are now reaching a point where small-effect loci are detectable, under the conditions that these variants are common : for instance, several GWAS studies of skin pigmentation levels have uncovered an amino-acid polymorphism of *MFSD12* as a determinant of colour variation across several continents (Adhikari et al., 2019; Crawford et al., 2017; Feng et al., 2021; Lona-Durazo et al., 2019). This gene is now a candidate that is also showing association signals in domestic animal studies (Hédan et al., 2019; Tanaka et al., 2019), suggesting it is effectively a hotspot of pigment variation. While human GWAS studies are systematically curated and will undoubtedly lead to powerful meta-analyses (Buniello et al., 2019), and while infrastructure is being built to integrate these data with laboratory model systems (Shefchek et al., 2020), there is a lack of resources to compile data from evolutionary and bred gene-to-trait relationships beyond these organisms. Gephebase, OMIA, and Animal QTLdb (Courtier-Orgogozo et al., 2020; Hu et al., 2021; Lenffer et al., 2006) are stopgap attempts at curating these data, but would need long-term resources to keep pace in front of increased rates of discovery in the genomic age.

## Conclusion

As more of the genetic variants underlying trait variation are identified, it becomes possible to more rigorously test predictions relating to the mechanisms of genetic evolution. Here, we highlight some of the trends in vertebrate pigmentation evolution, and specifically test some of the predictions made about the relative frequencies of *cis*-regulatory and coding mutations. In contrast to many people’s expectations, we found that the majority of the documented variation in pigmentation is driven by coding sequence mutations. However, we also identified multiple factors associated with mutational proportion that partly explain this disparity. We therefore made suggestions for the future direction of vertebrate pigmentation research with respect to both systems and study design. As the number and variety of case studies continues to increase, we expect our understanding of the genetic, cellular and developmental mechanisms underlying the evolution of vertebrate pigmentation to expand.

## Methods

### Literature curation in Gephebase

Gephebase compiles pairs of alleles in association with pairs of phenotypic states (a genetic variation causing a phenotypic variation is called a “gephe” for brevity). A full description of the database is provided in (Courtier-Orgogozo et al., 2020). In short, data currently cover all eukaryotes with relevant publications, with a focus on traits of evolutionary rather than clinical relevance. This includes variations that have been artificially selected by breeders (“Domesticated” dataset), or subject to experimental evolution under lab-controlled selective pressures. Data is manually curated by a team of less than a dozen researchers, using keywords as well as Pubmed/Google Scholar citations to identify newly published studies. Our current triage system gives priority to gephes identified by forward genetics (QTL mapping, GWAS) with a single-gene resolution and reasonable supporting evidence for causality (if not for the variant, at least for the gene). Gephes identified by candidate gene approaches without mapping are also included when there is additional functional evidence for the causality of the mutation. Gephebase is up-to-date and gathers all relevant papers published until 2017. Past 2017, data curation efforts have mainly focused on colour variation in vertebrates (Gephebase “Trait” category = “Coloration”). The download of Gephebase data on 28 October 2021 (Supplementary File 1), which was used for the present study, can be considered as a compilation of the genes and mutations contributing to Vertebrate coloration up to and including 2019.

### Meta-analyses

To examine trends in colour pattern evolution, we utilised Gephebase as a resource for exploring genotype-phenotype relationships. We formed a working dataset by selecting every Gephebase entry pertaining to the trait category ‘colouration’, and further filtered those entries by removing all entries pertaining to non-vertebrate species (accessed 28/10/2021). In total we retrieved 363 entries, with each entry representing a mutation at a single locus. Each mutation was present in only one gene for the organism in which it was identified, and there were a total of 61 unique genes identified across 89 vertebrate species. For each entry, we examined five parameters defined by Gephebase - Gene ID, Taxonomic Status, Study Methodology, Aberration Type, and Molecular Type (Courtier-Orgogozo et al., 2020). We further defined five additional parameters - Clade, Pigment Type, Phenotype Category, Functional Category, and Protein Category (Supplementary File 1).

### Assignment of new parameters

*Clade:* Each vertebrate represented in the dataset unambiguously belonged to one of five clades with distinct pigmentation biology - amphibians, teleosts, squamates, birds and mammals. As such, we divided entries into these clades to compare trends in their different pigmentation systems.

*Pigment Type:* In total, five pigment types were represented in the dataset - biliverdin, carotenoids, pteridines, melanins, and psittacofulvins. Additionally, one entry referred to structural coloration. Each entry could be unambiguously assigned to one or more of these categories. In total there were 10 entries in which a single genetic variant was associated with changes to multiple pigments. These entries were assigned to each relevant pigment type category.

*Phenotype Category:* We empirically assigned to each entry one of seven phenotype categories in order to group similar organismal phenotypes, and analyse trends in how they arise. Each entry was considered independently, and assigned the category that best describes the organismal phenotype based on the original study (Table 2). We accounted for both the visible phenotype in terms of pigment distribution and quantity, as well as the molecular mechanisms underlying the phenotype. For example, amelanism was distinguished from white spotting on the basis of whether the phenotype resulted from a lack of melanin synthesis, or a failure of melanocyte migration/differentiation. Ambiguous cases were not assigned a category - a total of 87 entries. This left a total of 276 assignments, of which 254 were *cis*-regulatory or coding. Additionally, categories were assigned relative to the ancestral state, so that for instance eumelanism refers to a mutation leading to an increase in eumelanin pigmentation.

We additionally assigned each category to either colour shift or pattern alteration, on the basis of whether the phenotype affected colour or spatial pattern boundaries. Each entry was considered independently, and these assignments were considered separately from the previous phenotype categories. Phenotypes involving a loss of patterning (for example whole body albinism resulting in no visible pattern boundaries) were considered colour shifts. Ambiguous entries were not assigned a category - a total of 76 entries. This left a total of 287 assignments, of which 263 were *cis*-regulatory or coding. This total (263) was used for the ‘total’ bars in Figures 4A, S3, and S4.

*Cellular Function Category:* In order to analyse genes associated with different developmental and cellular roles, we assigned each gene in the dataset a functional category, reflecting its role in the development of the phenotype (Supplementary File 1). Unlike phenotype categories, these assignments were based on the gene rather than on individual entries. Entries were assigned empirically, based on literature review of the genes identified by each study, and the phenotypic function(s) they have been previously implicated in. As with phenotype categories, each gene was assigned with high certainty and little ambiguity. Additionally, a total of four genes, comprising seven entries, were removed for the purposes of this analysis, as their cellular functions with regard to pigmentation biology specifically were unclear. These were *cyp19a1, copa, eif2s2, and lvrn*. Each gene had one entry each, except for *lvrn*, which had four entries.

*Molecular Function Category:* We assigned protein categories non-empirically by using the gene ontology (GO) terms associated with each gene in the dataset. We employed the EBI QuickGO Mouse Slimmer in order to identify parent GO terms associated with the sets of child terms associated with each gene. This slimmer was selected for being best representative of the distribution of GO terms within vertebrate clades. We then narrowed the set of GO terms to include only those associated with the ‘Molecular Function’ GO tree. We additionally manually combined a number of closely related GO terms in order to reduce the number of potential categories (Supplementary Table S1). In total there were 83 unique assignments of parent GO categories across the dataset. All of the 61 unique genes in the dataset generated at least one assignment, with the exception of *mlana*, a gene implicated in melanosome biogenesis. *Mlana* has only one molecular function assignment (GO:0005515 protein binding), which is not included in the QuickGO Mouse Slimmer. After combining closely related terms, there were 10 parent GO terms assigned. Then, we removed all categories with fewer than five unique gene assignments, of which there were three - RNA binding, enzyme regulator activity, and lipid binding. Thus we ended up with seven GO categories for comparison (Supplementary Table S1). The most commonly assigned GO category was the result of combining two GO terms - DNA binding and transcription factor activity. Although there are distinct biological differences between a gene displaying DNA binding activity and acting as a transcription factor, in the case of the Gephebase dataset there was nearly complete overlap between these terms. Only one gene was tagged as DNA binding without being tagged for transcription factor activity, namely egfr - epidermal growth factor receptor. All other genes were tagged with both GO terms, or neither.

*Evolutionary Hotspots*: We investigated the overlap of genes between pigmentation systems by examining the number of entries corresponding to shared genes. For each clade, we looked at all the genes identified in that clade, and then calculated the total number of entries corresponding to those genes in each of the other three clades. We then normalised each of these three figures by dividing by the total number of entries in that clade, in order to account for the disparity in entries between clades.

## Author contributions

JE performed all data analysis. JE, MES, AM and VC contributed to the study design. MES, AM and VC contributed to the gephebase curation. JE and MES wrote the manuscript with contributions or feedback from all authors. All authors read and approved the final version of the manuscript.

## Additional information

Supplementary File 1 contains the Gephebase dataset downloaded on 28 October 2021. It also contains information on parameter assignment (see methods) for each literature entry.

## Acknowledgments

We thank members of the Morphological Evolution Lab for discussion. AM and VC were supported by a John Templeton Foundation grant from 2014 to 2017 (JTF 43903). ES is supported by a NERC IRF NE/R01504X/1.

---

**Figure S1.**
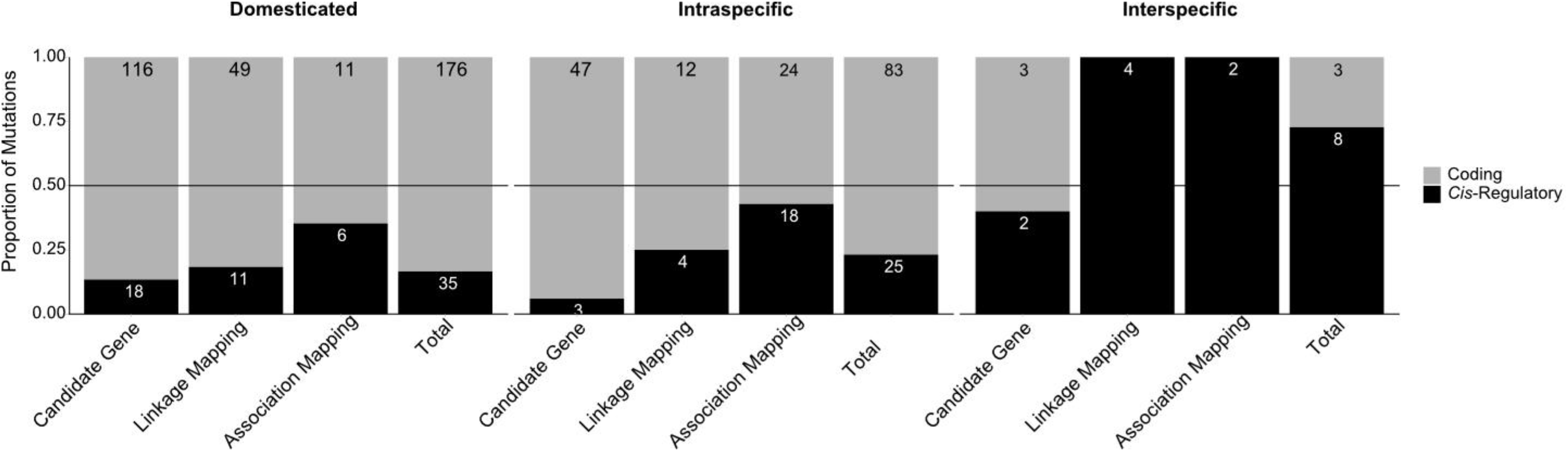
shows the proportion of *cis*-regulatory and coding mutations associated with different taxonomic statuses - domesticated, intraspecific and interspecific variation - controlling for study methodology, as well as the total proportion for each taxonomic status. The numbers above each bar represent the number of entries in that category. The grey horizontal line is at 0.5.

**Figure S2.**
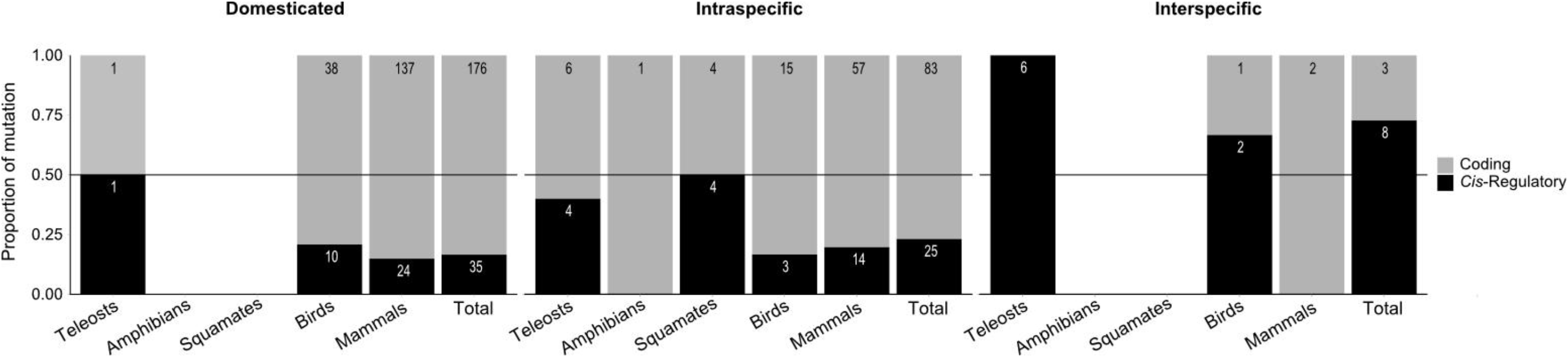
shows the proportion of *cis*-regulatory and coding mutations associated with different taxonomic statuses and clade - **A:** Domesticated variation, **B:** Intraspecific variation, and **C**: Interspecific variation. The total proportion for each taxonomic status is shown. The numbers above each bar represent the number of entries in that category. The grey horizontal line is at 0.5.

**Figure S3.**
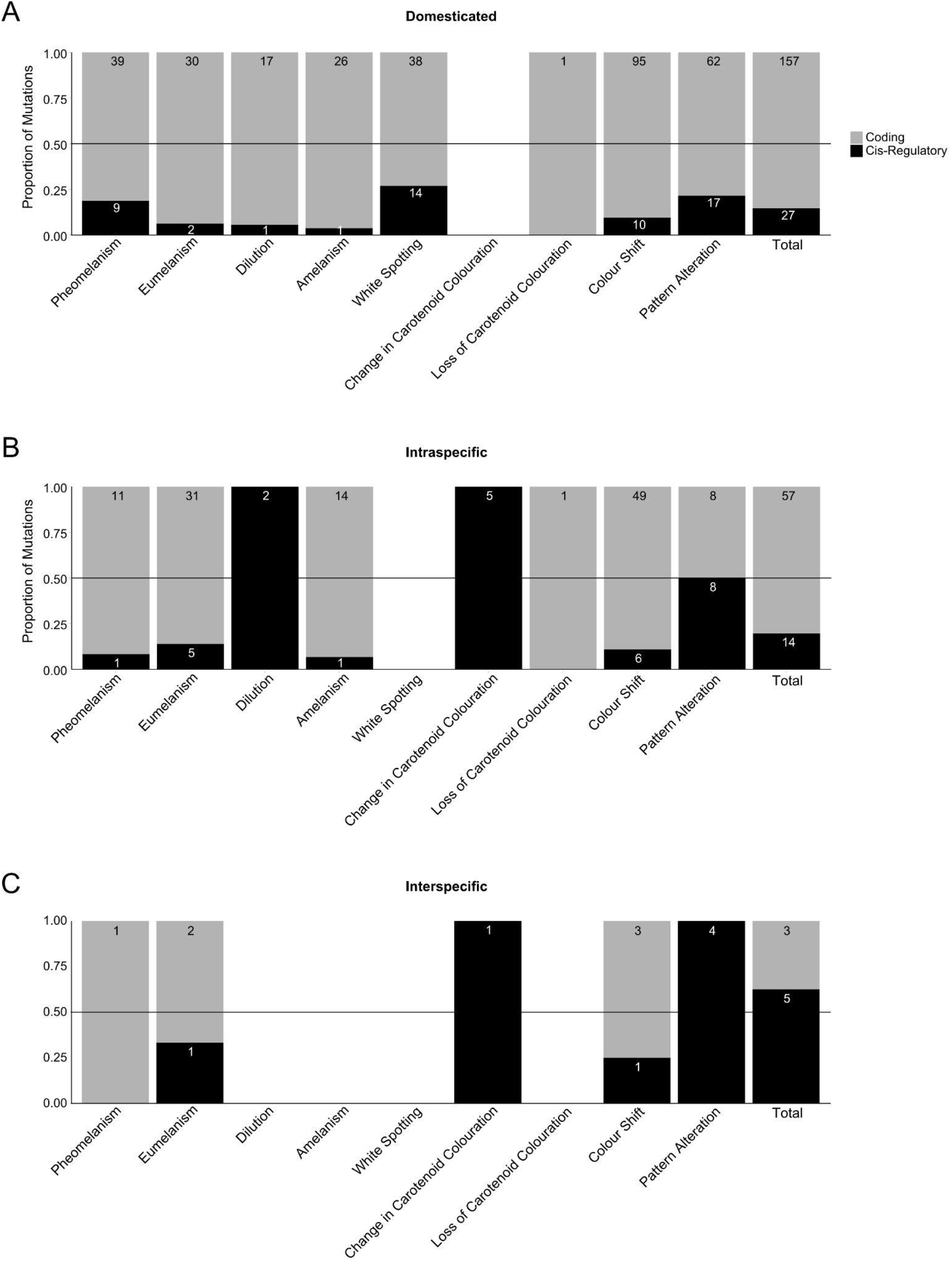
shows the proportion of *cis*-regulatory and coding mutations associated with different phenotype categories and taxonomic statuses - **A:** Domesticated variation, **B:** Intraspecific variation, and **C**: Interspecific variation. The total proportion for each taxonomic status is shown. The numbers above each bar represent the number of entries in that category. The grey horizontal line is at 0.5.

**Figure S4.**
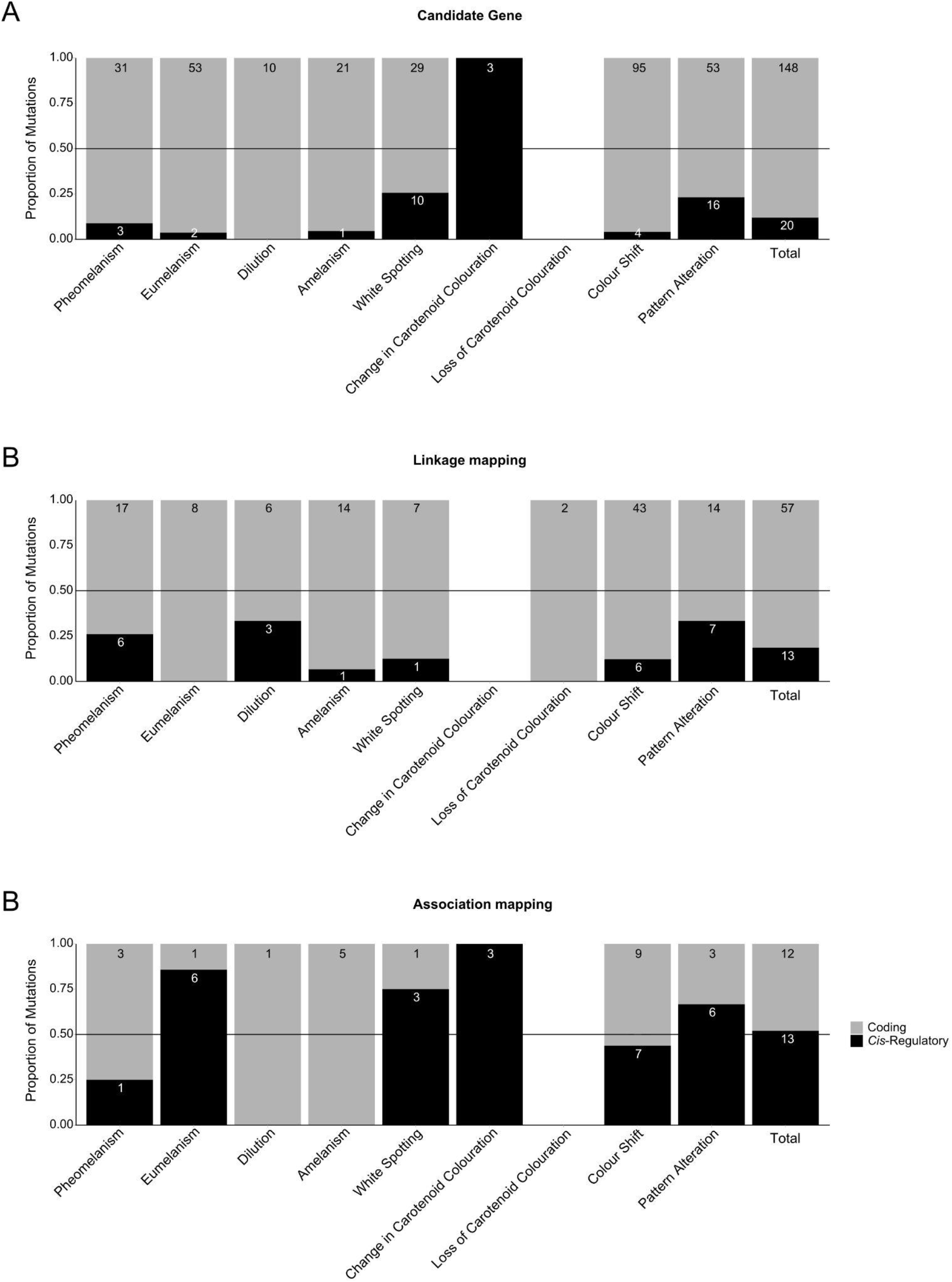
shows the proportion of *cis*-regulatory and coding mutations associated with different phenotype categories and study methodologies **A:** Candidate Gene, **B:** Linkage Mapping, and **C**: Association Mapping. The total proportion for each study methodology is shown. The numbers above each bar represent the number of entries in that category. The grey horizontal line is at 0.5.

**Figure S5.**
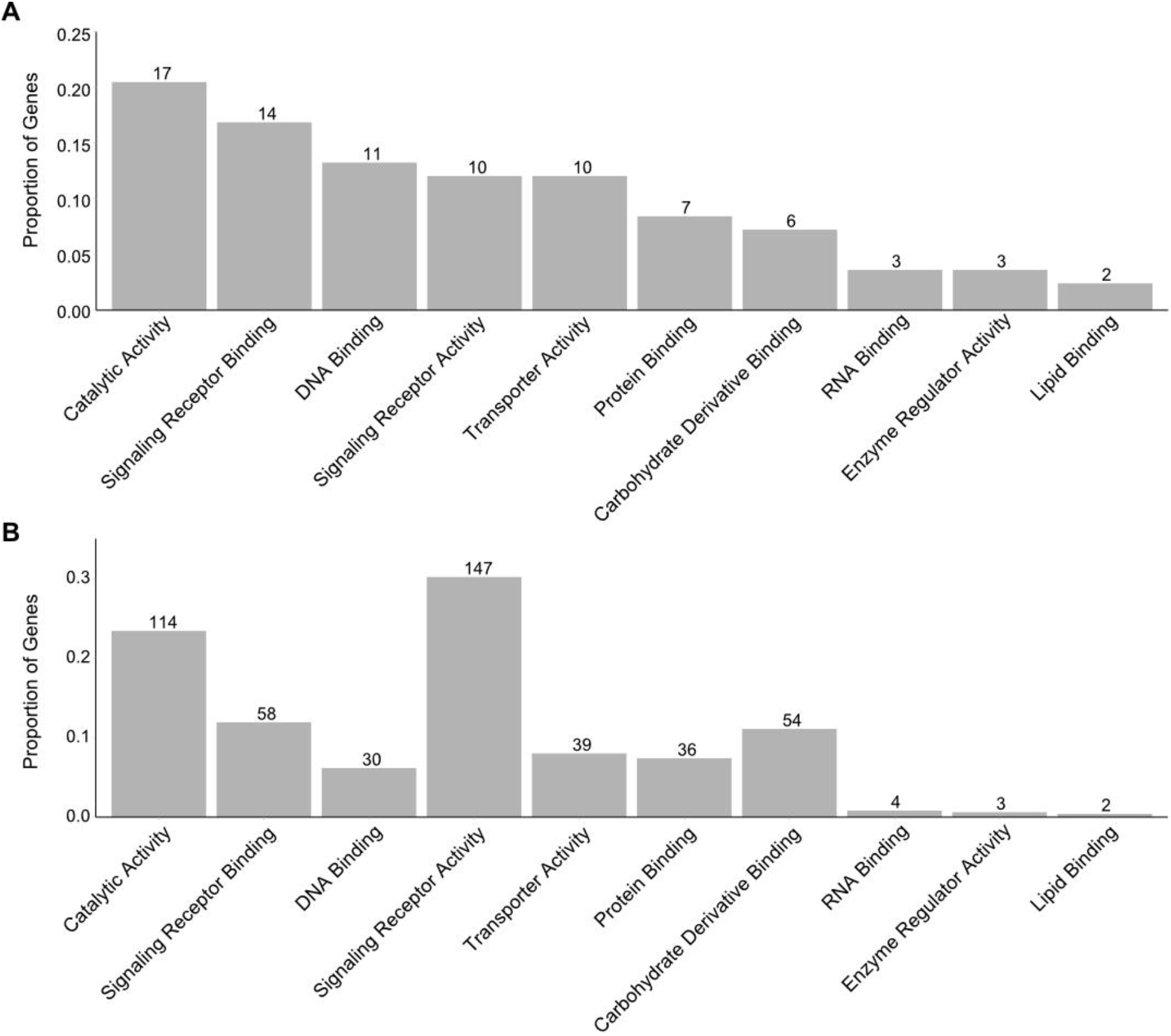
gene ontology assignments, unique genes and number of entries. **Figure S5, A:** The proportion of unique genes that each GO category was assigned to. **B:** The proportion of entries that each GO category was assigned to. The numbers above each bar represent the number of genes or entries in that category. Note that the RNA Binding, Enzyme Regulatory Activity, and Lipid Binding categories were removed for Figure 4C in the main text due to their low numbers of assigned genes (see Methods).

**Supplementary Table S1.**
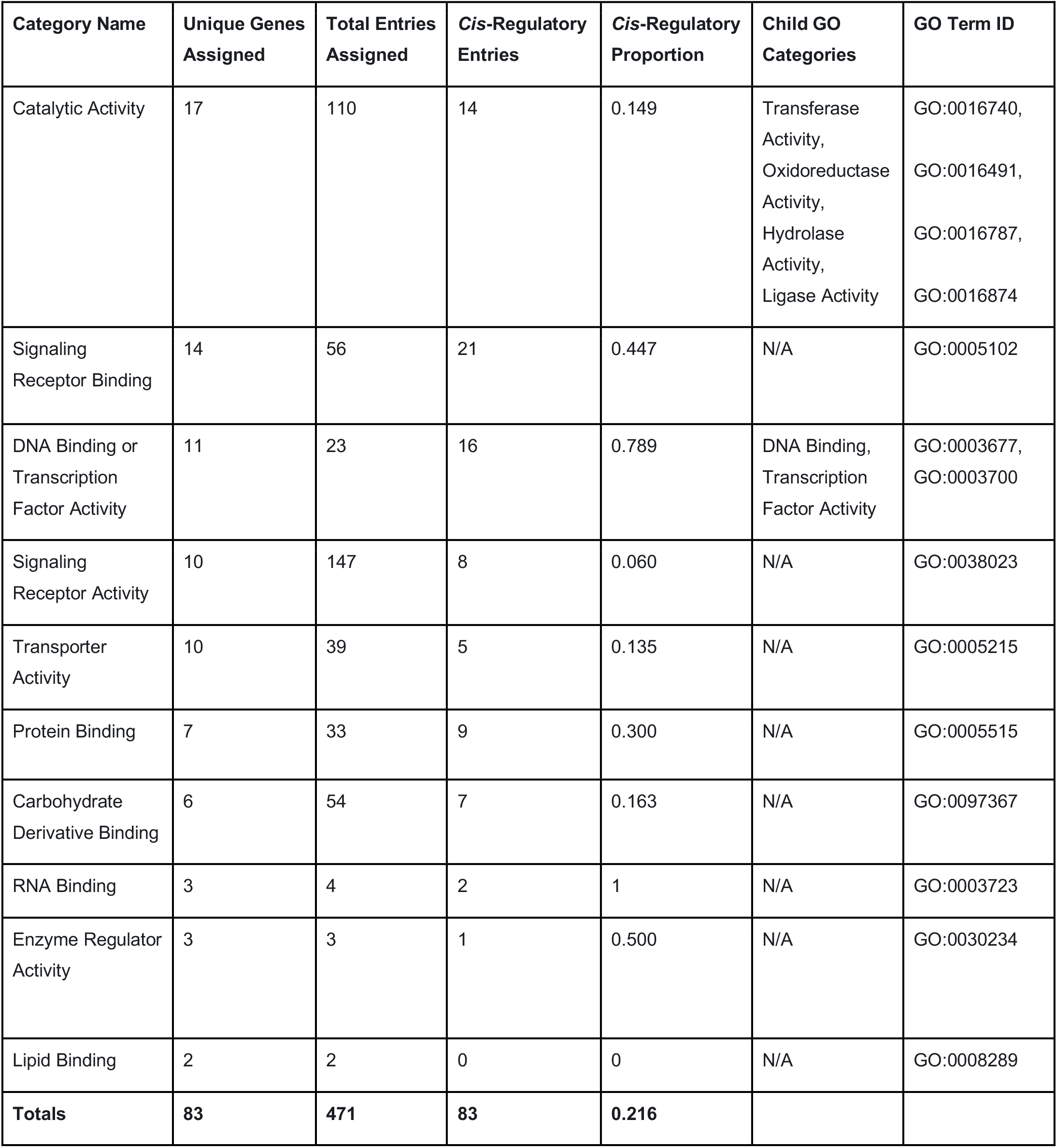
shows the 10 GO categories assigned to dataset entries, the total number of genes to which each GO category was assigned and corresponding total number of entries, the number of cis-regulatory entries to which each was assigned and corresponding cis-regulatory proportion, and the child GO terms that were combined into the category where relevant.

| Category Name | Unique Genes Assigned | Total Entries Assigned | Cis-Regulatory Entries | Cis-Regulatory Proportion | Child GO Categories | GO Term ID |
| --- | --- | --- | --- | --- | --- | --- |
| Catalytic Activity | 17 | 110 | 14 | 0.149 | Transferase Activity,<br>Oxidoreductase Activity,<br>Hydrolase Activity,<br>Ligase Activity | GO:0016740,<br>GO:0016491,<br>GO:0016787,<br>GO:0016874 |
| Signaling Receptor Binding | 14 | 56 | 21 | 0.447 | N/A | GO:0005102 |
| DNA Binding or Transcription Factor Activity | 11 | 23 | 16 | 0.789 | DNA Binding,<br>Transcription Factor Activity | GO:0003677,<br>GO:0003700 |
| Signaling Receptor Activity | 10 | 147 | 8 | 0.060 | N/A | GO:0038023 |
| Transporter Activity | 10 | 39 | 5 | 0.135 | N/A | GO:0005215 |
| Protein Binding | 7 | 33 | 9 | 0.300 | N/A | GO:0005515 |
| Carbohydrate Derivative Binding | 6 | 54 | 7 | 0.163 | N/A | GO:0097367 |
| RNA Binding | 3 | 4 | 2 | 1 | N/A | GO:0003723 |
| Enzyme Regulator Activity | 3 | 3 | 1 | 0.500 | N/A | GO:0030234 |
| Lipid Binding | 2 | 2 | 0 | 0 | N/A | GO:0008289 |
| **Totals** | **83** | **471** | **83** | **0.216** |  |  |

**Supplementary Table S2.** shows hotspot genes for each clade. For brevity, the 10 most represented genes for each clade are displayed, excluding genes with equal numbers of entries that would exceed this limit. For each gene, the number of entries in other clades is shown. Full data is in the provided Supplementary File 1.

| Gene | Number of<br>Mammal Entries | Number of Bird<br>Entries | Number of<br>Teleost Entries | Number of<br>Squamate Entries |
| --- | --- | --- | --- | --- |
| **Mammals** |  |  |  |  |
| MC1R | 59 | 24 | 2 | 3 |
| Kit | 49 | 0 | 0 | 0 |
| Asip | 29 | 6 | 0 | 0 |
| Tyr | 18 | 7 | 0 | 0 |
| Tyrp1 | 14 | 2 | 0 | 0 |
| SLC45A2 | 11 | 6 | 2 | 0 |
| PMEL | 11 | 3 | 0 | 0 |
| Mlph | 7 | 2 | 0 | 0 |
| **Birds** |  |  |  |  |
| MC1R | 59 | 24 | 2 | 3 |
| Tyr | 18 | 7 | 0 | 0 |
| Asip | 11 | 6 | 2 | 0 |
| SLC45A2 | 29 | 6 | 0 | 0 |
| PMEL | 11 | 3 | 0 | 0 |
| Ndp | 0 | 3 | 0 | 0 |
| Sox10 | 0 | 3 | 0 | 0 |
| **Teleosts** |  |  |  |  |
| Pax7 | 0 | 0 | 4 | 0 |
| Oca2 | 2 | 0 | 3 | 1 |
| MC1R | 59 | 24 | 2 | 3 |
| SLC45A2 | 11 | 6 | 2 | 0 |
| **Squamates** |  |  |  |  |
| MC1R | 59 | 24 | 2 | 3 |
| Oca2 | 2 | 0 | 3 | 1 |
| BCO2 | 0 | 2 | 0 | 1 |
| PREP | 0 | 0 | 0 | 1 |
| Prkar1a | 0 | 0 | 0 | 1 |
| Spr | 0 | 0 | 0 | 1 |

